# 4D Single-Particle Tracking with Asynchronous Read-Out SPAD-Array Detector

**DOI:** 10.1101/2023.08.25.554867

**Authors:** Andrea Bucci, Giorgio Tortarolo, Marcus Oliver Held, Luca Bega, Eleonora Perego, Francesco Castagnetti, Irene Bozzoni, Eli Slenders, Giuseppe Vicidomini

**Affiliations:** Molecular Microscopy and Spectroscopy, Istituto Italiano di Tecnologia, Genoa, Italy; Dipartimento di Informatica, Bioingegneria, Robotica e Ingegneria dei Sistemi, University of Genoa, Genoa, Italy; Non coding RNAs in Physiology and Pathology, Istituto Italiano di Tecnologia, Genoa, Italy

**Keywords:** single-particle tracking, fluorescence lifetime, confocal laser-scanning microscopy

## Abstract

Single-particle tracking (SPT) techniques are essential for investigating the complex functions and interactions of individual, specifically labelled particles in biological environments. Many SPT techniques exist, each optimised towards a different balance between spatiotemporal resolution and range, technical complexity, and information content. This bargain is exemplified by the contrast between wide-field camera-based and real-time SPT approaches, with the latter being generally more advanced but at the cost of high complexity. Furthermore, the fluorescence lifetime, a powerful tool for investigating the particle’s interactions and nano-environment, has yet to be measured consistently.

To overcome these limitations, we propose a novel real-time three-dimensional SPT technique based on a hybrid approach. In our implementation, we equip a confocal laser-scanning microscope with an asynchronous read-out single-photon avalanche diode (SPAD) array detector and few other optics. Each sensitive detector element acts as a confocal pinhole, and the recorded intensity distribution reflects the particle’s position in three dimensions relative to the excitation volume. This localization is used in a real-time feedback system to keep the particle in the centre of the excitation volume. Importantly, as each pixel is an independent single-photon detector, SPT is combined with fluorescence lifetime measurement.

Our system achieves a localization precision of up to 30 nm with 100 photons and microsecond time resolution, while also performing fluorescence lifetime measurements. First, we validated the technique by tracking fluorescent particles in artificial environments. Secondly, as further validation, we investigated the movement of lysosomes in living SK-N-BE cells and measured the fluorescence lifetime of the GFP marker expressed on a membrane protein. We observed an unprecedented correlation between the changes in fluorescence lifetime and the motion state of the lysosomes.

Thanks to its simplicity and the great momentum of confocal microscopy based on SPAD array detector, we expect that this implementation will open to many information-rich correlative imaging and tracking studies.

## Main

Fluorescence single-particle tracking (SPT) is a fundamental tool to investigate the complex functions, structures, and interactions of individual particles in biological environments [1]. As an example, it has been successfully applied to study virus infection mechanisms [2–4], surface protein trafficking [5–8], molecular motors dynamics [9, 10] and anomalous diffusion and transport [11, 12].

The high variability of biological questions, which can be addressed by SPT, has pushed the method development in different directions. Although many approaches have been proposed, no one has emerged as the gold standard.

Off-line SPT is the first and most straightforward method: the sample is illuminated in a wide-field configuration, and the fluorescence emission is recorded with a megapixel matrix detector such as a complementary metal oxide semiconductor (CMOS) or charge-coupled device (CCD) camera. Repeated exposures over time produce a set of images that are analyzed off-line to extract the positions of the single particles, while the trajectories are reconstructed by linking the localizations frame by frame [8, 13–15]. This approach is capable of following many particles in parallel with a spatial resolution between 20 nm and 40 nm given a budget of 10^3^-10^4^ photons [8, 16, 17], but is severely limited to ≈10 ms in time resolution by the imaging rate of the detector and to 2 µm in axial range due to the fixed illumination plane [17].

An alternative class of techniques is real-time single-particle tracking (RT-SPT). This approach aims to overcome the temporal resolution and spatial range limitations of off-line SPT. Real-time SPT retrieves the position of only a single particle inside a small observation volume and follows its movement over time by shifting the observation volume via a closed feedback loop. Common RT-SPT techniques are implemented on customized laser-scanning microscopes featuring one or more single-pixel detectors, such as single-photon avalanche diodes (SPADs) or photo-multiplier tubes (PMTs). Therefore, experimental raw data consists of one (or more) intensity time traces in which the spatial information is encoded using structured detection or structured illumination. In the first approach, the emission is split among multiple detectors, whose spatial arrangement is engineered to allow the inverse calculation of the 3D particle’s position from the intensity traces [18–20]. Structured illumination, on the contrary, investigates the particle’s location by sequentially moving the focused excitation beam at a certain number of points along a specified trajectory around the particle. The ideas proposed across the years are diverse and include the use of a Gaussian beam swiping a circle, such as in orbital tracking [21–23], or illuminating points arranged in a tetrahedron [24] or moved along a knight’s tour [25]. More recently, in MINFLUX a doughnut-shaped beam is displaced in a triangular pattern, showing a more efficient localisation precision for a given number of photons. [26–29]. Indeed, given a certain photon budget, the uncertainty of any RT-SPT technique can be theoretically calculated depending on the combination of the illumination shape and the spatial arrangement of the sampling points [30].

Regardless of the actual implementations, all RT-SPT techniques are affected by one or more of the following issues: high photon fluxes requirement, poor axial range, and technical complexity. Furthermore, the information about the photon arrival time is often not exploited, despite being available in many cases. This prevents, for example, access to the fluorescence lifetime measurement, which is a powerful tool for investigating the particle’s interactions and chemical nano-environment [31, 32]. A more detailed discussion about the state of the art of RT-SPT can be found in the comprehensive review by van Heerden et al. [33].

In recent years, companies, as well as research groups, have directed growing attention toward the realization of better detectors. The development of new cameras (such as Hamamatsu qCMOS, Photonscore LINcam, and event-based sensors [34, 35]) aims to improve the temporal resolution and information content of wide-field microscopy, including off-line SPT. In the same way, advances in SPAD technology, and in particular the development of asynchronous read-out SPAD arrays, are beneficial for laser scanning microscopy techniques [36, 37]. An asynchronous read-out SPAD array detector is constituted by a set of independent SPADs placed on the same chip with a predetermined spatial arrangement. Each pixel works as a standalone detector with single-photon detection and timing capabilities. It is directly wired to its output channel, which fires a digital signal with high (a few tens of picoseconds) temporal precision for each detected photon. In such a fashion, the detector can be combined with a multi-channel time-resolved data-acquisition system (e.g., a photon time-tagging module) to provide spatial information like a small camera while overcoming the frame-rate limitation. Indeed, by having a completely pixel-based asynchronous read-out, the camera’s frame rate is theoretically infinite. Only the so-called pixel dead time practically limit the frame-rate: after detecting a photon, the pixel is blind for a few tens of nanoseconds. Additionally, every photon can be tagged with its arrival time with respect to the excitation laser pulse, thus giving access to the fluorescence lifetime information. SPAD arrays detector have already been employed to increase the resolution and contrast of confocal images through image scanning microscopy [38–43] and to enhance the information content and flexibility of fluorescence correlation spectroscopy [44]. Both imaging and spectroscopy approaches have been synergistically combined with fluorescence lifetime analysis [45].

Here, we present a novel feedback-based RT-SPT implementation based on a common laser scanning microscope equipped with a 5×5 asynchronous read-out SPAD array detector and an astigmatic detection. The detection scheme provides direct and almost instantaneous particle’s 3D localization across a relatively small static field-of-view (sFoV) – approximately the size of the focal region, i.e., sub-micrometers in all directions (*x, y, z*). With this information we can dynamically re-position the beam scanning system to maintain the particle centered in the static field-of-view (sFoV). Consequently, the effective FoV of the RT-SPT system extends to cover the complete scanning range of the microscope, spanning hundreds of micrometers in all directions. Furthermore, the SPAD array detector simultaneously measures the particle’s fluorescence lifetime *τ* .

As such, our RT-SPT implementation effectively produces a 4D trajectory in time and space [*x*(*t*)*, y*(*t*)*, z*(*t*)*, τ* (*t*)], thus improving the information content while reducing the architectural complexity – in comparison to any approach mentioned above. We call this implementation real-time 4D single-particle tracking (RT-4D-SPT). Here, there is no need to perform orbital scanning or other fast excitation beam shifting to encode the single particle’s position. Notably, the localisation of the particle – for the sFoV re-positioning – is calculated in real-time by the data-acquisition (DAQ) system, without introducing a sensitive delay in the feedback closed loop system. The spatiotemporal resolution of the proposed RT-4D-SPT approach is practically limited only by the flux of the photon detected and by the lag-time of the laser beam scanning system – to re-centering onto the particle before it escapes from the sFoV. This later condition can however be relaxed by enlarging the sFoV. In addition, the full SPAD array signal is transferred to the computer for refining the particle trajectory off-line. We validate our technique by comparing the localization uncertainty obtained from the Cramér-Rao bound calculations with the experimental localizations of fixed fluorescent particles. We then track fluorescent particles in two configurations: moved along a predetermined path and freely diffusing in a viscous solution. To prove the versatility of our technique, we apply it in a biological context to investigate the movement of lysosomes in living cells along microtubule filaments while assessing the lifetime of the green fluorescent protein (GFP) marker expressed on a protein on their membrane.

## Results

### Principle of RT-4D-SPT

To perform RT-4D-SPT, we use a 3D laser scanning microscope with minimal modifications (Fig. 1a). The fluorescence emission generated from the Gaussian excitation region at the scanning position **r**_s_ = (*x*_s_*, y*_s_*, z*_s_) is made astigmatic by a cylindrical lens and focused onto the SPAD array detector located into a conjugate plane – and which replaces the traditional confocal pinhole and the single-pixel detector. Here it is essential to highlight that the SPAD array detector registers the fluorescent signal in de-scanned mode, thus the probed region (i.e., excitation region) and the sFoV are always co-aligned. Our SPAD array detector is a specialized device that combines the temporal performance of a SPAD with the structured detection of a 5×5 camera, imaging the small (1.4 A U) sFoV in the sample. When a photon is detected by a sensitive element/pixel of the array at coordinates (*i, j*), it produces a short electric pulse in the corresponding output channel, with a time jitter below 200 ps. The signal is fed into a multi-channel time-resolved data-acquisition (DAQ) system, which allows for measuring the photon detection time *t*_d_. Thus, each detection event corresponds to an independent 6-dimensional data point (*x*_s_*, y*_s_*, z*_s_*, i, j, t*_d_).

**Fig. 1.**
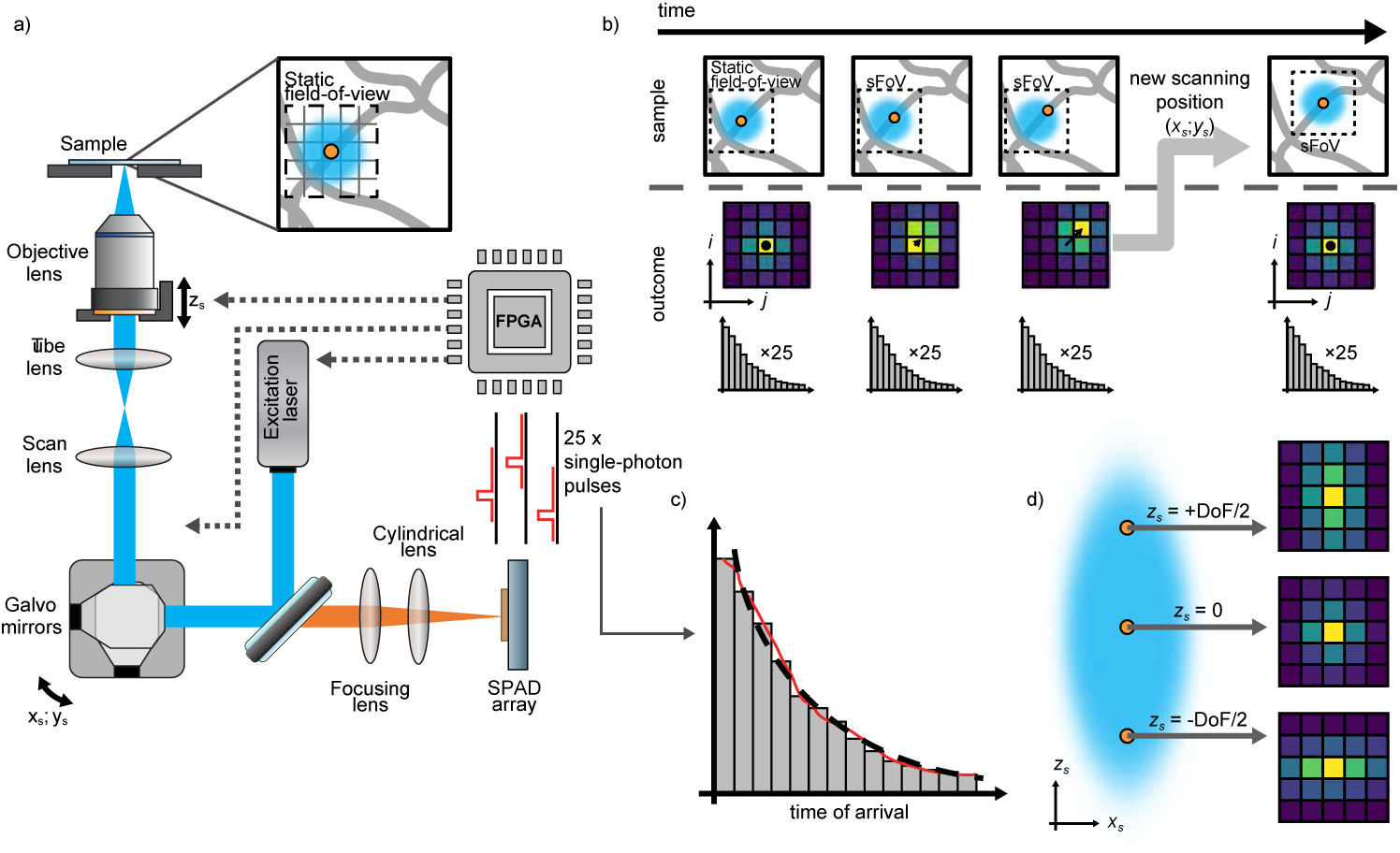
real-time 4D single-particle tracking on a laser-scanning microscope equipped with a SPAD array detector. **a)** The optical setup is based on a 3D laser scanning microscope by replacing the confocalized detection unit with a SPAD array detector and inserting a cylindrical lens. The detector static field-of-view (≈1.4 A U) is composed, in our case, of 5×5 elements leading to 25 independent single-photon pulse trains. The field-programmable gate array (FPGA) receives the trains, calculates the particle position and updates the 3D beam position in real-time. **b)** A 2D movement of the illuminated single particle with respect to the center of the excitation results in a shift of the detected emission pattern and a reduction in its intensity (as shown in the microimages). The new position (*x_s_, y_s_*) can therefore be estimated and the sFoV is re-centered onto the particle with the beam positioners. **c)** The temporal performance of each SPAD element of the array detector is similar to the single element counterpart (time jitter *<*200 ps). Thus, the fluorescence lifetime of the emitter *τ* can be extracted from the histogram of the delays between the photon detection events and the pulsed excitations. **d)** The quantitative estimation of the axial position of the particle leverages the astigmatism caused by the cylindrical lens. In fact, the symmetry and shape of the emission pattern registered in the microimage changes throughout the depth-of-field (DoF).

To obtain the emitter’s position in real-time, we can use position estimators developed for wide field microscopy, such as the centroid estimator. The centroid estimator needs low computational costs and can be easily implemented directly in the field programmable gate array (FPGA) high-speed DAQ board. This results in a fast – potentially down to 100 ns – feedback which is leveraged for the sFoV re-positioning. In detail, the centroid estimator analyzes the 5×5 images *I*(*i, j*), referred to as *microimages*, obtained by accumulating the detected photons stream within a specific and tunable temporal window. Any particle movement produces a distinguishable change in the intensity distribution in the microimage. Specifically, a lateral displacement leads to a shift in the distribution’s centre while, by making the detection astigmatic, its shape is uniquely linked to the axial position [46–48] (Fig. 1d). The same DAQ board also acts as a control unit for the whole microscope and its beam scanning apparatus: A pair of galvanometer mirrors for the lateral displacement and a piezo objective for the axial shift – for our implementation. The control unit delivers the input signals to the galvanometer mirrors and objective piezo to displace the sFoV in the new position.

In short, our method for real-time tracking utilizes the spatio-temporal information provided by the SPAD array detector to repeatedly re-center the sFoV onto the particle (Fig. 1b). Because of the single-photon asynchronous readout of the detector, the re-centering can be decided *a-priori* at a specific frequency, i.e., every fixed time interval Δ*t*_rc_ – as for synchronous detector with limited frame rate – or, more interestingly, when a specific event occurs. For example, when the number of photons detected reaches a particular value, namely when a suitable signal-to-noise ratio is obtained.

Whilst the sFoV reaches only a few hundred nanometers, when using high numerical aperture objective lens, the effective achievable tracking range can easily exceed hundreds of micrometres, depending on the dynamical range of the lateral and axial positioners and the field number of the objective lens. Regarding spatiotemporal resolution, the size of the sFoV plays an important role. Indeed, the re-centering action is necessary only to prevent the particle from escaping from the sFoV. Larger is the sFoV, and smaller are the temporal constraints for the re-positioning. Furthermore, tracking the particle can also be performed within the sFoV without re-centering. As a result, the achievable tracking rate is higher and decoupled from the update rate of the scanning position (*x_s_, y_s_, z_s_*).

Because the information collected by the SPAD array detector is also transferred to the personal computer (PC) in the form of microimages with a high frame-rate, the spatiotemporal precision of the tracking can be improved off-line using a more precise and robust localisation algorithm than the centroid.

Not less important than deciphering the particle’s position, it is to study the particle’s fluorescence lifetime as a function of time. By using a pulsed excitation laser and implementing a series of fast time-to-digital converters (TDCs) directly in the DAQ system, we can obtain the delay between the excitation event and the photon detection, the so-called photon arrival time, Δ*t*_em_ = *t*_d_ − *t*_exc_. This information enables us to calculate the particle’s fluorescence lifetime *τ* – potentially in real-time (Fig. 1c). In this work, we implemented within the FPGA-based DAQ system 25 TDCs by using a digital frequency domain (DFD) architecture. With respect to more precise TDC implementation based on tapped-delay lines, the DFD architecture requires less FPGA resources and sustains the maximum photon-flux achieved by the SPAD array detector. Specifically, our DFD implementation provides a sampling step (or temporal resolution) of 397 ps, a value lower than the tens of picoseconds obtained by using TDC based on tapped-delay lines, but still optimal for fluorescence lifetime application. The low FPGA-resources allow to synergistically integrate the multiple TDC in the same DAQ module implementing the tracking and controlling the microscope.

### Localization within the sFoV

In RT-SPT, the ability to track a single moving particle hinges on the precise, accurate, and timely localization of its position. This enables the re-centering of the probed region – and thus the detection volume – on the particle before the particle leaves the detection volume. Consequently, the characterization of localization performance stands as a critical first step in thoroughly assessing any tracking methodology. Here, we characterise the localisation precision within the sFoV for our RT-4D-SPT approach.

According to estimation theory, one way to quantify the theoretical lower limit of the localization uncertainty is to calculate the so-called Cramér-Rao bound (CRB) [49]. The CRB is a statistical measure that provides a theoretical lower bound on the variance of any unbiased estimator of an unknown parameter. In our case, the unknown parameter is the position of the particle being tracked.

To calculate the CRB, we adapt the mathematical framework developed by Masullo et al. for sequential structure illumination single-molecule localisation microscopy [30]. Specifically, we introduce some minor modifications to account for our structured detection scheme (Supplementary Note SI Note 1:). Our calculation is founded on three key assumptions: (1) both the signal and background photon counts follow a Poisson distribution, (2) the signal is linearly dependent on the excitation light intensity, and (3) the background is independent of the excitation intensity and scanning position. These assumptions are typically valid in low-excitation experiments, where fluorescence is far from saturation and the primary background source is the detector’s dark noise. Considering a fixed acquisition time, the total number of photons detected by the sensor, denoted as *N* (**r**_e_), and consequently the signal-to-background ratio (*SBR*(**r**_e_)), are dependent on the position **r**_e_ of the particle relative to the center of the sFoV. However, the experimental localization conditions can be fully characterized by the scalar parameters *N*_p_ = *N* (**0**) and *SBR*_p_ = *SBR*(**0**). Henceforth, these benchmarks will be extensively employed.

We based the CRB calculations on the experimental PSFs measured by using 20 nm fluorescent beads. By assuming *N*_p_ = 100 photons and *SBR*_p_ = 5, we obtain the maps of the CRB in the lateral 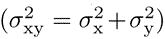 and axial 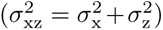 planes (Figs. 2a-f). From the isoline curves we can identify a volume of roughly 300 nm 300 nm 500 nm in which we expect an approximately flat localization uncertainty with a maximum value between 40 nm and 60 nm.

**Fig. 2.**
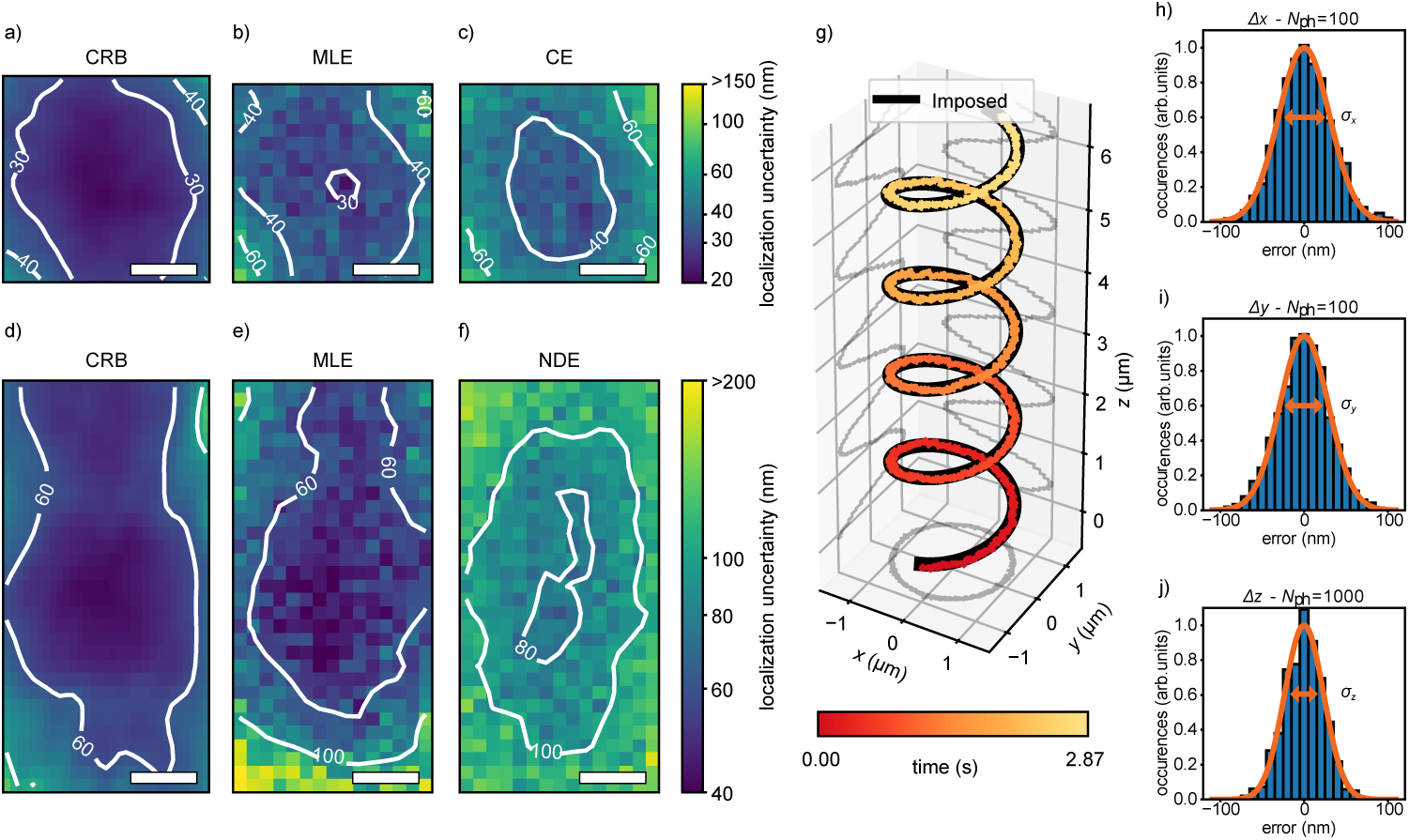
Characterization of the planar localization and tracking uncertainties. **a-c)** Lat-eral uncertainty maps *σ*_xy_(*x*_e_, *y*_e_). The Cramér-Rao bound (CRB) is calculated using an experimental point spread function (PSF), *SBR*_p_ = 5 and *N*_p_ = 100 photons and assuming a fixed acquistion time. The uncertainty maps of the maximum likelihood estimation (MLE) and centroid estimator are mea-sured by using a common dataset obtained by scanning 20 times a 20 nm fluorescent bead replicating the same conditions of the CRB. Scalebar = 100 nm. **d-f)** Axial uncertainty maps *σ*_xz_(*x*_e_*, z*_e_). The CRB is calculated using an experimental PSF, *SBR*_p_ = 5 and *N*_p_ = 100 photons and assuming a fixed acquistion time. The uncertainty maps of the MLE and normalized difference estimator are mea-sured by using a common dataset obtained by scanning 33 times a 20 nm fluorescent bead replicating the same conditions of the CRB. Scalebar = 100 nm. **g)** Imposed and tracked trajectories of a single 20 nm fluorescent bead (*λ*_exc_ = 561 nm) moved at 10 µm*/*s along an helical pattern. The scanning position is updated laterally with the centroid estimator every 100 photons 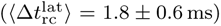 and axially with the normalized difference estimator every 1000 photons 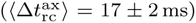. The color is used to visualize time. **h-j)** Histograms of the localization uncertainty along the three direc-tions for the experiment in **a**. *σ*_x_ = 30.5 ± 0.3 nm, *σ*_y_ = 29.0 ± 0.4 nm, and *σ*_z_ = 23.2 ± 1.2 nm.

Amongst all the localization estimators proposed by different techniques through the years, the MLE has emerged for its reliability and solid mathematical background [46, 50, 51]. Remarkably, it provides an unbiased and linear estimation (Supplementary Note SI Note 2:) and it is demonstrated to be fully efficient, which means it asymptotically reaches the CRB.

The high computational complexity necessary to perform the estimation with the MLE makes it the ideal choice for off-line analysis but hinders its implementation in the real-time loop. We consequently need a new set of faster estimators, which allows us to gain computational speed at the expense of accuracy. We therefore introduce the centroid and the normalized difference estimators for the lateral and axial localizations, respectively. Compared to the MLE estimator both the centroid and the normalized difference are faster but less resistant to noise and their linearity is affected by the signal-to-noise ratio (SBR). Furthermore, the normalized difference estimator for the axial localization is not independent from the lateral position of the emitter (Supplementary Note SI Note 2:).

To compare the planar localization uncertainty of each estimator with the theoretical lower limit, we localise a single 20 nm fluorescent bead replicating the emitting conditions of the CRB calculations. Using the 3D piezo stage, the particle is shifted across the entire sFoV and localized multiple times with both the MLE and the faster estimators at each position. We then take the standard deviation as a measure of the localization uncertainty to plot the lateral (Figs. 2b,c) and axial (Figs. 2e,f) planes for the MLE (Figs. 2b,e), the centroid (Fig. 2c), and the normalised difference (Fig. 2f) estimators.

The comparison of the MLE results with the theoretical lower limit shows a slightly worse performance in the global minimum value. This can be justified by the additional noise sources that may occur during the experiment but are not included in the CRB model, such as sample and microscope drifts, light scattering and other sources of background. As expected, the performance with the faster estimators further deteriorates, especially with the normalized difference estimator.

By looking at the lateral and axial uncertainty line profiles (SUpplementary Fig. S1), we observe the overall shape of the uncertainty maps is nevertheless consistent with the CRB. We define the optimal detection volume as the region in which the uncertainty increases at most 50 % with respect to the global minimum. Hence, we identify a volume of at least 300 nm 300 nm 600 nm inside which the performance of the localization is considered optimal with all the estimators. In particular, the faster estimators provide a lateral planar uncertainty between 35 nm and *≈*53 nm and an axial planar uncertainty between 80 nm and *≈*120 nm.

Considering a minimum effective re-centering time Δ*t*_rc_ 1 ms, we can predict this optimal detection volume provides a maximum measurable diffusion coefficient for Brownian motion of 5 10 µm^2^*/*s. This upper limit matches with the diffusion coefficients of relatively small proteins (above 30 − 40 kDa) moving in the cytoplasm and the membrane of cells [52–54], while it may be unsuitable for diffusion in pure water [55] – depending on the size of the particle.

### Real-time 3D and 4D tracking

The tracking procedure leverages the localization information by updating in real-time the beam scanning position (*x*_s_*, y*_s_*, z*_s_) to keep the single particle always in focus. To test the validity of our approach in a controlled scenario, we first track a fixed 20 nm fluorescent bead moved along a 3D helical path at 10 µm*/*s (Fig.2g). As already anticipated, due to timing constraints, the emitter position **r**_e_ has to be retrieved with the centroid and the normalized difference estimators, which are less precise than the MLE and potentially non linear.

To evaluate the precision of our tracking system and check whether these drawbacks affects the performance of the feedback loop, we compute the histograms of the difference between the imposed and tracked trajectories in three dimensions (Figs. 2h-j) and fit them with a gaussian function. Assuming the scaling factor 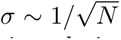, we can therefore calculate the lateral and axial planar tracking uncertainties relative to the detection of 100 photons *σ*_xy_ = 42.0 0.4 nm and *σ*_xz_ = 79.5 3.4 nm. We compare this outcome with the planar localization uncertainties of the fast estimators and we observe the values obtained from the tracking experiment are compatible with the scenario in which we localize a particle placed in the central region of the sFoV. Remarkably, the localization uncertainty obtained during tracking is not sensibly worse, despite the potential presence of additional sources of noise such as the positioners jitter and inertia. Therefore, the dominant source of uncertainty entering the feedback loop is still the estimation process itself. Furthermore, the experiment also suggests that the feedback loop is correctly implemented and well calibrated so that it effectively converges fast enough to retain the particle inside the sensitive region.

The 3D tracking capabilities of the system are now tested in a less ideal scenario by following freely diffusing fluorescent beads. In this experiment, four different water-glycerol dilutions characterized by the volumetric ratio *R*_gly_ = *V*_gly_*/V*_tot_ are created, and 40 nm beads are added at a concentration sufficiently low so that the average number of particles inside the sFoV at any time is less than one. For each dilution *R*gly we then collect a set of *N*_tr_ independent trajectories **r**^(^**^n^**^)^(*t*) = **r**^(^**^n^**^)^(*t*) from which we calculate the MSD averaging over all the different trajectories and time intervals as described in 4. Each MSD curve (Fig. 3a) exhibits a behaviour that is compatible with the hypothesis of 3D Brownian motion described by MSD(*δt*) = 6*D*_c_*δt*, which is expected for particles suspended in a homogeneous liquid [56]. By linearly fitting the graphs with the previous formula, we can calculate the average diffusion coefficient *D*_c_(*R*_gly_) and subsequently the average hydrodynamic radius *r*_hyd_ applying the Stokes-Einstein equation. The four measurements provide an average bead diameter of 32 13 nm, which is compatible with the value 37 6 nm reported by the seller, further proving the effectiveness of our approach.

**Fig. 3.**
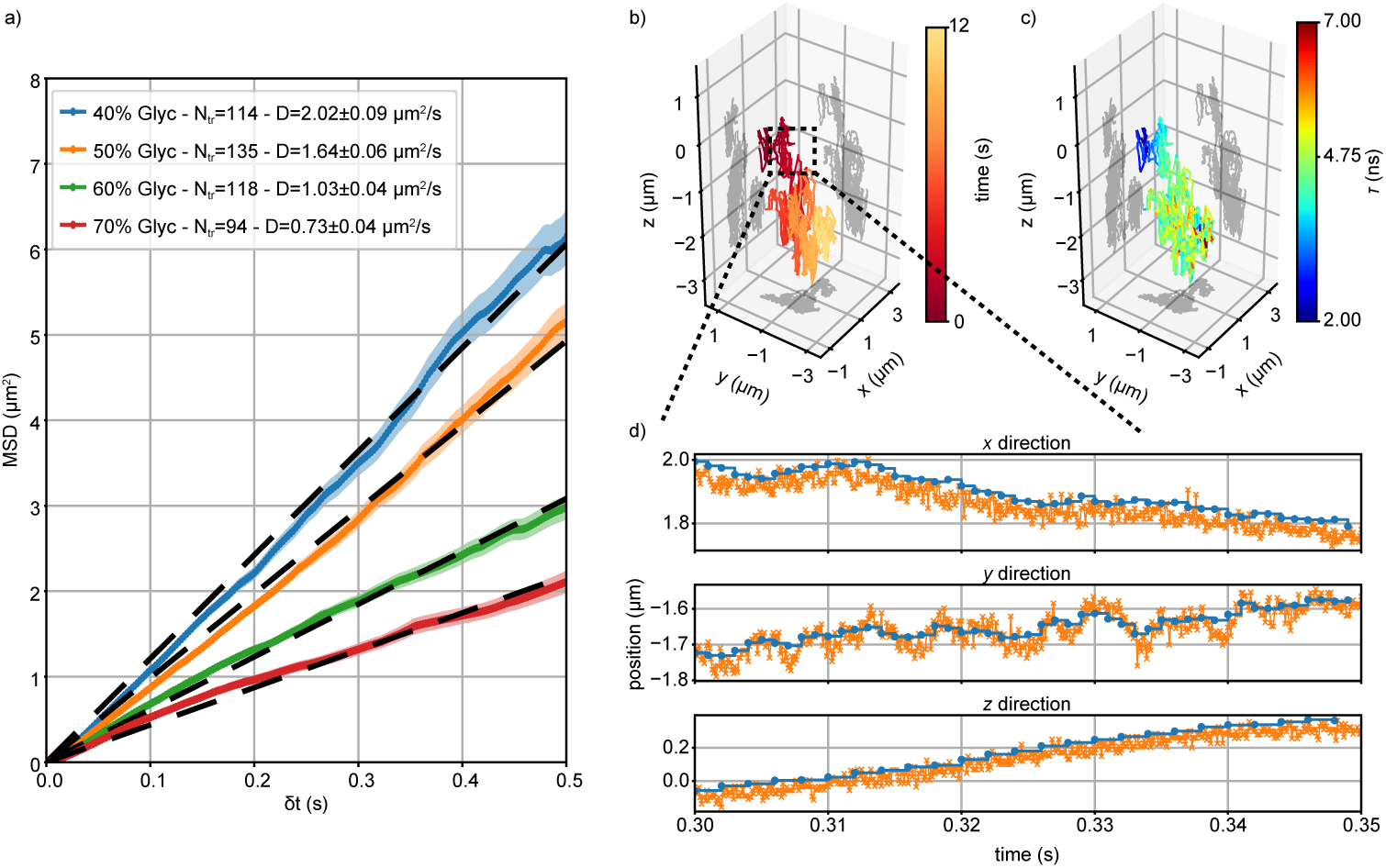
3D and 4D tracking on free fluorescent beads. **a)** Mean squared displacement (MSD) of 40 nm fluorescent beads (*λ*_exc_ = 561 nm) diffusing in different glycerol/water volumetric dilutions *R*_gly_. Each curve is averaged over a number *N*_tr_ of independent trajectories obtained with real-time tracking at a fixed re-centering time 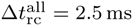. Under the assumption of 3D Brownian motion, the MSD is linearly fit to extract the corresponding diffusion coefficient *D*_c_(*R*_gly_). **b)** 3D representation of the trajectory of a single 100 nm fluorescent bead (*λ*_exc_ = 488 nm) diffusing in a *R*_gly_ = 70 % glycerol-water solution. The color visualizes the elapsed time. The re-centering in the lateral and axial directions is performed at 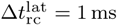 (⟨*N* ⟩ = 582 photons) and 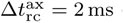 (⟨*N* ⟩ = 1164 photons) respectively. **c)** 3D representation of the same trajectory as **b**, but with color used to visualize the fluorescence lifetime *τ* . The estimation is performed every Δ*t_τ_* = 1 ms. **d)** Spatial trajectory rebinned in postprocessing with a dwell time of 100 µs in all the directions. The localization is also refined by using the MLE.

Our capabilities are not limited to 3D spatial tracking alone. We can also simultaneously register the evolution of the fluorescence lifetime *τ* (*t*). The FPGA control module manages the synchronization of the pulsed excitation and photon detection, allowing for the native computation in real-time of the 25 histograms of the photon emission time Δ*t*_em_. To achieve this, we implement 25 parallel TDCs, one for each SPAD pixel, utilizing the DFD principle [57, 58]. Specifically, with our implementation, we can obtain a complete histogram of the emission time with a sampling step of 397 ps every 5.7 micros (Tortarolo et al., submitted).

To show the richness of our real-time output, we perform *4D tracking* by measuring the 4-dimensional trajectory Trj(*t*) = [*x*(*t*)*, y*(*t*)*, z*(*t*)*, τ* (*t*)] of a 100 nm fluorescent bead freely diffusing in a *R*_gly_ = 70 % glycerol-water solution. Such a dataset cannot be visualized in a single graph and requires slicing along a subset of the available dimensions depending on the phenomenon being studied. A first intuitive representation focuses on the spatial information by plotting the 3D trajectory in space and adding an additional observable through color coding (Figs. 3b,c). We notice the fluorescence lifetime seems to exhibit a space-dependent behaviour with a region near the beginning of the trajectory associated to a value of around 1.5 ns, which then fades to 5 ns as the particle diffuses. However, these observations based solely on the spatial information do not provide a solid explanation of the phenomenon. In fact, in this specific case study, a deeper understanding is obtained with a different representation of the 4D trajectory, where the observable are analyzed as a function of time. By calculating the MSD curve (Fig. S2a) we deduce the particle is undergoing anomalous subdiffusion MSD(*δt*) ∝ *δt^α^* with *α <* 1. While this result appears to contradict previous measurements of beads diffusing in the same dilution (Fig. 3a), the subdiffusion regime could be caused by spatial heterogeneity in the solution [59] due to the incomplete mixing of water and glycerol in this particular experiment. Furthermore, the time evolution of the fluorescence lifetime (Fig. S2b) reveals the presence of self-quenching – which causes a decrease in lifetime when a high amount of fluorophores are packed in proximity to each others, i.e. in a big fluorescent bead. As the emitters bleach (hypothesis confirmed by the progressive decrease in the fluorescence photon-flux), this condition relaxes and the fluorescence lifetime gradually reaches a value associated with the fluorophore in isolated conditions . A reduction of the detected photons during the fixed estimation window Δ*t_τ_* = 1 ms also leads to an increase in noise.

Notably, this single experiment demonstrates the benefits of having a 4D dataset, which allows us to simultaneously reveal both spatial properties such as the diffusion regime and spectral properties, for example, self-quenching.

### Off-line tracking and postprocessing

Our SPT technique belongs to the macro area of the real-time approaches, but the information content of the data is richer than its usage in real-time and eventually emerges with postprocessing analysis, as typically happens in off-line tracking. As a consequence, it is not possible to define a single value for the temporal resolution and the characterization of the speed of the system requires a more detailed discussion.

Firstly, the photon-countings data in their digital form are generated in the FPGA control module at a reading rate of 508 MHz. Therefore, the lowest time resolution of any possible digital mechanism is ≈2 ns.

Processing the photon stream to produce a re-centering response is a task that requires a non-negligible amount of computational time. The processing pipeline is described in details in the materials and methods supplementary section 4. We experimentally measured that a triggering input on the digital card is translated into an actuated voltage output of the analogue card with a time delay of 400.0 2.5 ns. Nev-ertheless, the actuation is further slowed down by the setting time of the positioner, requiring ≈300 µs for the galvanometric mirrors (lateral shift) and ≈10 ms for the objective piezoelectric stage (axial shift). The overall real-time procedure is therefore limited to a response time of around 1 ms, which means any faster phenomena will generally produce an averaged trajectory, if any.

However, the effective time resolution of our technique is decoupled from the maximum trackable speed as our approach features an off-line postprocessing part fed with highly resolved data (Fig. S3). In fact, the re-centering action is necessary only to keep the particle inside the sFoV and, under this condition, the localization itself could be performed at any integer multiple of the 2 ns digital quantization time. For this reason, the control module sends the microimages and the scanning position to the PC every 2 µs and the lifetime histogram in a tunable integer multiple of 10 ns. The dataset resulting from a valid trajectory is therefore natively temporally resolved in the microsecond regime.

As an example, in Fig.3d we disply a portion of the trajectory depicted in Fig.3b that has been rebinned at 100 µs and whose localization estimation is refined with the MLE estimator. The postprocessing pipeline not only adds a 10-fold (lateral) and 20-fold (axial) improvement in time sampling but also reveals previously hidden spatial fast movements that were averaged out in real-time due to the longer integration window.

### Motion of lysosomes in living cells

Lysosomes are membrane-enclosed organelles that play a critical role in cellular homeostasis by removing damaged organelles, as well as extraneous particles through phagocytosis [60]. Dysfunction of lysosomes has been linked to a range of diseases, including lysosomal storage disorders [61, 62], neurodegenerative diseases [63, 64], and cancer [65–67]. Tracking their movement is therefore a crucial task as it can help developing new strategies for treatments or prevention of lysosomal-related disorders [68–70]. Of particular interest are the lysosomes found at the cell periphery which show high dynamicity [71, 72]. Their motion pattern is characterized by stationary states, when the lysosome is bound in one location (“stop” states), alternated with fast movements between locations (“run” states) [72–74].

To study this “stop-and-run” alternating behaviour, we fluorescently labeled the lysosomial membrane with GFP by tagging the lysosome-associated membrane protein LAMP-1. We therefore performed 4D tracking on the organelles found at the periphery of living human neuroblastoma SK-N-BE cells. The cellular context is provided by a reference bi-dimensional (2D) image (*x, y*) of the microtubule structure acquired before any tracking experiment (Fig. 4a and supplementary Video SV1). We observe that the region of the sample explored by the lysosome has a diameter of approximately 7 µm, which considerably exceeds the dimension of the sFoV. As expected, the lysosome appears to move between adjacent sites, following quasi-rectilinear paths, whose shapes agree well with the microtubule structure determined by imaging. The discrepancies between the trajectory and the microtubules structure is mostly attributed to the different dimensional of the two datasets, i.e., the trajectory spams across a 3D space, while the image is a single 2D optical section.

**Fig. 4.**
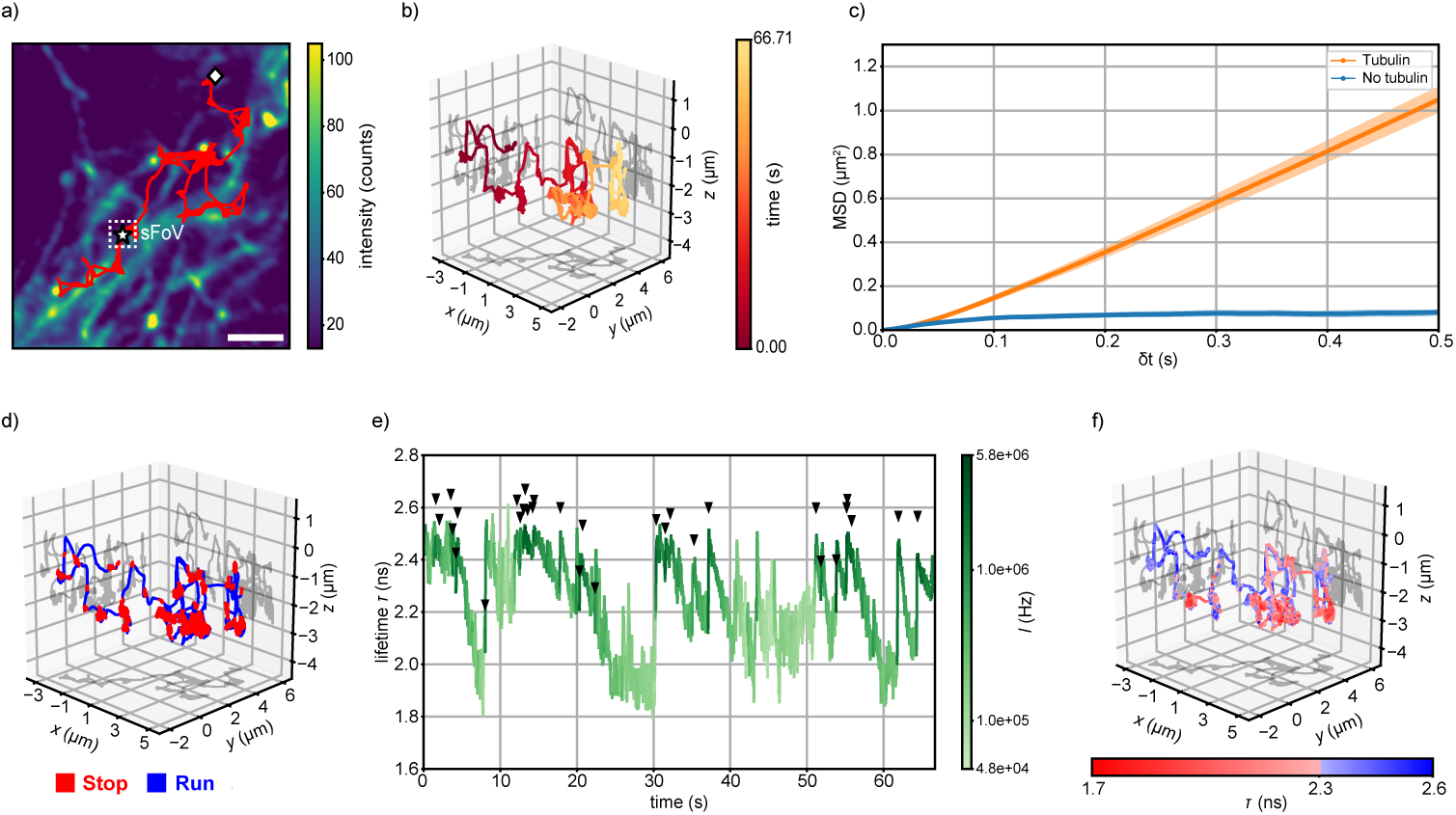
Investigation of lysosomes diffusion with RT-4D-SPT. **a)** 2D projection of the spatial trajectory of a single lysosome moving inside a living neuroblastoma SK-N-BE cell. A white star and a white romb indicate the beginning and the ending of the trajectory respectively. The organelle is tracked by exciting the GFP expressed on a membrane protein (*λ*_exc_ = 488 nm) with the re-centering performed at a fixed timing 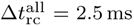. The reference image shows the tubulin proteins labeled with Abberior LIVE 560 (*λ*_exc_ = 561 nm) and is acquired prior to the lysosome tracking measurement. Pixel dwell time of 100 µs. Scalebar = 2 µm. **b)** 3D plot of the trajectory shown in **a**. The colormap represents the temporal scale. **c)** Average mean squared displacement of lysosomes moving in living neuroblastoma SK-N-BE cell before and after the addition of nocodazole (*n* = 15). Both curves are obtained by averaging the MSD of 15 independent lysosome trajectories 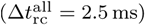. The average duration of the tracking is 53 ± 25 s in presence of tubulin and 5 ± 4 s after nocodazole addition. **d)** 3D plot of the trajectory shown in **a**. The color represents the diffusion state of the lysosome. Segmentation is performed by thresholding the average time spent by the lysosome inside voxels of dimension 5 nm × 5 nm × 20 nm. **e)** Time evolution of the fluorescence lifetime *τ* for the trajectory in **a** (Δ*t_τ_* = 10 ms). The color represents the fluorescence intensity. The black arrows indicate the peaks where the photon flux increases of at least 1 MHz with respect to the previous value. **f)** 3D plot of the same trajectory of **a** with the color indicating the fluorescence lifetime value.

To provide additional evidence of the role of the microtubules, we compare the motility of lysosomes before and after the addition of nocodazole, a drug that interferes with microtubule polymerization [75, 76]. To do so, we compute the average MSD of different independent trajectories acquired before and after the addition of the drug (Fig. 4c). The de-polymerization of microtubules clearly impacts the movement of the lysosomes, which consequently exhibit a strongly subdiffusive behavior, indicating a local confinement in space (supplementary Fig. S4).

To unveil the “stop-and-run” motion pattern of the lysosomes, we developed a simple yet effective quantitative segmentation procedure. Briefly, the two states are differentiated by computing the residence time of the lysosome in neighbouring positions inside the cell, with the “stop” states identified therefore by long residence times. This approach is able to reveal the dual motion of the lysosomes in 3D (Fig. 4d).

When considering the full 4D dataset provided by the experiment, we can additionally analyze the time trace of the fluorescence lifetime (Fig. 4e). We measure a not-constant value of the fluorescence lifetime during the motion of the lysosome. Specifically, the fluorescence lifetime value gradually drops from the expected value for the GFP *τ* ≈ 2.5 ns [77] to a plateau at 1.9 ns. The peak value is suddenly recovered on average every 3.4 ± 0.5 s. Interestingly, the fluorescence photon-flux varies as well between peak values of approximately 3 6 MHz and minima as low as 10 kHz. Remarkably, the 4D trajectory reveals a strong correlation between the two processes (Fig. 4e). By adding the fluorescence lifetime information on a 3D plot of the trajectory (Fig. 4f ans Supplementary Video SV2), we observe the value of the fluorescence lifetime correlates with the motion states of the lysosome. Specifically, values around 2.4 ns are associated with the “run” state, while the gradual drop below this value is associated with the “stop” state.

In the proposed case study of single lysosomes tracking, the information about fluorescence lifetime decay greatly simplifies and improves the analysis of the lysosome motion state. This behavior is confirmed to a broader extent in the Supplementary Note, where we collect a pool of 15 independent 4D lysosome tracking experiments and analyze them with an automated segmentation algorithm based only on the fluorescence lifetime, hence not requiring any user intervention.

## Discussion

The unique nature of asynchronous read-out SPAD array detectors merges important abilities of off-line and real-time SPTs. This class of sensors works as a camera detector – with a small sFoV and virtual sub-microsecond frame-rate – and as a SPAD detector – with single-photon sensitivity and timing. By combining the SPAD array detector with a fast FPGA-based data-acquisition module, the virtual photon-counting images are used to compute in real-time the position of the particle. This position represents the input of a feedback close loop system which maintains the single particle of interest in focus – within the sFoV – by driving the laser beam scanning microscope apparatus. An off-line procedure can then post-process the same virtual images for refining and rebinning the trajectory with a time resolution multiple of 2 µs. Because the real-time feedback system must simply maintain the particle within the sFoV, whilst a precise particle localisation can be obtained off-line, the FPGA-based card can implement a fast but less accurate particle position estimation. As a result, the proposed SPT approach’s primary limitation is the actuators’ lag time for re-positioning the microscope’s focus. For this reason, we envisage a series of new *RT* 4*D SPT* implementations in which the focus’ and static field-of-view’s sizes are changed according to the apparent diffusion coefficient of the particle. Indeed, the excitation region and sFoV can be tuned by controlling the laser beam size and the overall microscope magnification on the detector plane, respectively. Notably, the proposed combination of real-time and off-line localisation is also compatible with those techniques which sequentially move the beam around the particle to estimate its position (e.g., MINFLUX and orbital tracking).

A fundamental characteristic of the proposed RT-SPT approach is the ability to measure the fluorescence lifetime, opening truly *RT* 4*D SPT* experiments. The particle can be followed across a large 3D effective field-of-view, while the fluorescence lifetime can be used to understand the particle’s interactions or changes in its nano-environment. Because we implemented the measurement of the fluorescence lifetime on the same DAQ and control module used for real-time particle tracking, this approach can open to smart experiments in which some actions (e.g., the activation of a second laser beam) are triggered by a fluorescence lifetime change.

To demonstrate the importance of correlating dynamics information with the fluorescence lifetime, we conducted live cell experiments which monitor the diffusion behaviour of lysosomes. Our findings revealed an interesting correlation between the intermittent “stop-and-run” motion pattern and the intensity and the fluorescence life-time of the GFP expressed on the organelles’ membrane. While the drop and recovery in intensity can possibly be explained with the physical 3D rolling of the organelle in the two alternating states [78] – which could expose new fluorophores on the top surface – the variation in the fluorescence lifetime of the GFP is an unprecedented observation. Without further investigations, it isn’t easy to provide a valid explanation for this phenomenon. However, it is worth mentioning that the GFP is sensitive to many parameters of its environment, including refractive index [79] and pH [80]. Thus, the change in fluorescence lifetime could signal the organelle’s membrane rearrangement during its function.

In conclusion, our ability to measure both the position and the fluorescence lifetime of a particle in real-time using a single instrument significantly broadens the range of biological phenomena that can be observed. Our RT-4D-SPT technique, which combines simplicity and informativeness, can potentially set a new gold standard. With the rapid development of SPAD array technologies, our approach represents a promising avenue also for single molecule tracking experiments, where the photon-flux value substantially reduces.

## Methods

### Optical setup details

The backbone of the optical setup for the proposed RT-SPT method is a conventional laser scanning microscope in which we replace the typical single-element detector with a 5×5 asynchronous read-out CMOS SPAD array detector placed in a conjugate image plane of the microscope (Fig. S5).

Laser light at the wavelength *λ*_exc_ = 561 nm (MPB VFL-P-1000-560) and at *λ*_exc_ = 488 nm (PicoQuant LDH-D-C-485) is combined and modulated in amplitude with an acousto-optic modulator (AA Opto-Electronic MT80-A1-VIS) in a dedicated laser box. Using a single-mode polarization-maintaining fiber (Thorlabs P5-405BPM-FC-2) the light is brought to the microscope setup, collimated with a reflective collimator (Thorlabs RC08FC-P01) and reflected towards the objective at a multi-color dichroic beam splitter (Semrock Di01-R405/488/561/635-25×36). The beam passes through a first telescope formed by lenses L2 (LINOS G063-232-000) and L1 (LINOS G063-237-000) and it is scanned in the lateral directions (*x*_s_ and *y*_s_) via a pair of galvanometric mirrors (Thorlabs GVSM002-EC/M). A second telescope composed by the scan lens (Thorlabs SL50-CLS2) and the tube lens (Thorlabs TTL200MP) magnifies the collimated beam to an effective diameter of ≈46 mm, further cropped by the usage of 1 inch optical elements. Nevertheless, the back-aperture of the 63x/1.40 oil objective (Leica HC PL APO 63x/1,40 OIL CS2) is overfilled. Finally, the objective lens focuses the excitation beam into the sample. The axial position of the focal point *z*_s_ is adjusted by moving the objective along the axial direction with a linear piezoelectric stage (Physik Instrumente PIFOC P-725.2CL). The sample is moved in the micrometer scale with a manual 3D microstage (Piezoconcept Manual Microstage Nikon + Piezoconcept Rectangular manual Z adjust) and controlled in the nanometer scale with a 3D piezoelectric stage (Piezoconcept BIO3.300) placed onto of the microstage. The emitted fluorescence is collected in epifluorescence mode by the same objective lens, de-scanned by the galvanometric mirrors and transmitted at the dichroic beam splitter into the detection arm. The remaining leakages of the laser illumination are blocked with two notch filters (Chroma ZET488NF and Chroma ZET561NF). The fluorescence light passes through the cylindrical lens (Thorlabs LJ1516RM-A) to induce astigmatism and it is finally focused onto the SPAD array detector by lens L3 (LINOS G063-238-000). The setup is contained in a relatively small breadboard of 800 mm × 800 mm (Standa 1B-A-80-80-015-BL) and features an overall theoretical magnification from sample plane to the SPAD image plane of *M* = 504.

### Control module

All the actions and computations necessary to control the microscope during the tracking experiments are performed in real-time by the dedicated control module. The control module is governed by a custom firmware developed in LabVIEW, which is integrated with a Graphical User Interface (GUI) on the PC (the “host”). This setup allows for dynamic user intervention and real-time data visualization. The design includes a chassis (NI PXIe-1071) housing a thunderbolt-based communication module (NI PXIe-8301) and two FPGA-based data-acquisition cards (the “targets”), operating in a master/slave configuration (Supplementary Fig. S6). Specifically, the chassis facilitates synchronization among all components through a shared clock running at 10 MHz, and enables internal communication between the two acquisition cards via eight independent trigger lines. We employ an I/O digital board (NI PXIe-7822R) to handle the signals from the SPAD array, and an I/O analog and digital board (NI PXIe-7856R) for interfacing with the positioners and laser sources.

The digital card serves as the system’s central hub by collecting the single-photon pulses from the SPAD array and implementing the real-time feedback loop. The digital input from each SPAD element is checked at regular intervals of ≈1.97 ns = 1*/*508 MHz, ensuring lossless sampling of the readout regardless of the selected SPAD array hold-off time. The single-photon signals are then accumulated in the incremental photon-counting registers **n**. To measure the photon timing, the TDC unit processes the photon-counting registers using the DFD approach, enabling the calculation of a full histogram of the detection times **t**_d_ every 5.7 µs = 1*/*176 kHz. Both **n** and **t**_d_ are then fed to the logical unit, where they are processed at a cycle clock frequency of 20 MHz to trigger a real-time response.

Each new scanning position **r**_s_ generated by the logical unit is immediately transmitted to the analogue card with a custom-written protocol that utilizes the eight internal trigger lines of the chassis. This protocol allows an entire message to be deliv-ered in under 190.0 2.5 ns. The analogue card, acting as a slave, blindly converts control orders from external sources into voltage outputs. While the scanning position is determined by the digital card as explained above, the piezoelectric stage position and the lasers power are delivered from the host PC and can therefore be directly modified by the user. The speed of the electronic actuation is fixed by the digital-to-analog converter (DAC)’s minimum update time of 1 µs = 1*/*1 MHz, as specified by the seller.

The experimental raw data consists in the full status of the control system and it is sent to the host PC at two distinct timings. The photon-countings registers **n** and the scanning position **r**_s_ are buffered at intervals of 2 µs = 1*/*500 kHz, while the time of detection histograms **t**_d_ are buffered at *k* 10 ns = *k/*100 MHz, where *k* is a tunable integer used to adjust the data rate and manage communication bandwidth.

The effectiveness of the feedback loop in locking the sFoV onto a single moving particle mainly depends on the logical unit. The current implementation only uses the photon-counting information **n** to check whether a new scanning position **r**_s_ is required. At the beginning of each experiment, the user sets the specific behaviour of the logical unit, including the update conditions for each spatial axis and the choice of localization estimators. Indeed, the logical unit allows for independent refreshing of the lateral (*x*_s_*, y*_s_) and axial (*z*_s_) scanning positions. In particular, the photon countings **n** are utilized to evaluate three main conditions: the presence of a particle inside the detection volume, the need for lateral re-centering, and the need for axial re-centering. The algorithm is structured into four main steps (Supplementary Fig. S7): collection, evaluation, decision and re-centering. Each step is executed at the cycle clock frequency of the logical unit. After the initialization of the environment, in the “collection” step the photon-counting registers **n** are read and utilized to update the internal variables integrating the number of detected photons since the last lateral (*N*_xy_) and axial (*N*_z_) re-centering events. In addition, the algorithm also keeps trace of the total amount of detected photons (*N*_found_) and the elapsed time (Δ*t*_found_) since the last determination of the presence of a particle.

In the subsequent “evaluation” step, the algorithm checks whether the user-defined conditions for the re-centering are satisfied, either in the lateral or axial direction. The user can choose between two alternative strategies: in the *photon-counting* mode, the evaluation is positive when the integrated registered photon-countings *N*_xy_ or *N_z_* reach a specified threshold value, whereas in the *fixed dwell time* mode the triggering takes place at a predetermined rate, regardless of the number of photons. In case it is needed, new hybrid modalities which consider the value of the fluorescence lifetime can easily be added.

When either the lateral or axial directions requires a re-centering, the scanning position does not immediately change. Instead, the system goes through the “decision” step, during which the update request is reviewed to ensure the photons are coming from a particle. To discriminate between a proper signal and the background, the system checks the values of *N*_found_ and Δ*t*_found_ against a user-defined minimum signal photon flux. If the elapsed time exceeds the maximum waiting time without registering at least a minimum threshold amount of photons, then the data is labeled as background and all the variables are reset. Consequently, the re-centering request is also discarded.

If the review is positive – the signal is considered valid – the process finally proceeds to the “re-centering” step, where a new scanning position is calculated.

### Mean squared displacement calculation

Considering a set of *N*_tr_ independent trajectories **r**^(**n**)^(*t*) = **r**^(**n**)^(*t*) of length *L*^(*n*)^ sampled with a discrete sampling step *dt*_s_, we calculate the MSD as follows:

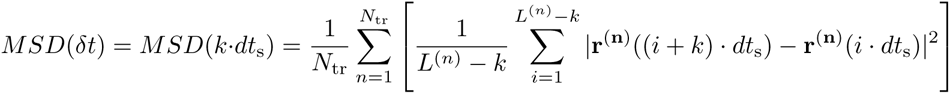

It is important to note that the calculation averages over all the different trajectories and time intervals, implicitly assuming the ergodic hypothesis for the single-particle trajectories [81]. Furthermore, this formulation utilizes all the available displacements of duration *k dt*_s_, increasing the averaging pool for each MSD point, but at the same time exposes to the risk of correlations between overlapping displacements.

### Cell line and sample preparation

#### Fluorescent beads

To produce a sample with fixed fluorescent beads, we first deposit 150 µL of poly-L-lysine (PLL) on a clean cover slip and incubate it at 37 *^◦^*C for 10 min. Meanwhile, we prepare a dilution of the beads’ mother solution in distilled water with a volumetric ratio between 1:500 and 1:1000, which we sonicate for 5 min. We dry the coverslip with clean air and deposit 150 µL of beads dilution on top of the adhesive film, followed by incubation at 37 *^◦^*C for 10 min. We spill the remainder of the solution on the cover glass and dry it with clean air. We add 5 µL of Mowiol^®^ mounting medium and seal the cover glass on a microscope slide. A summary of all the beads used in the various experiments can be found in Tab. ST1

#### Live cells

SK-N-BE neuroblastoma cells line are cultured in RPMI medium 1640 (Gibco), supplemented with 10 % fetal bovine serum (FBS), GlutaMAX (Gibco), and penicillin/streptomycin, and induced to differentiate by 10 µM all-trans-Retinoic acid (RA, Sigma) for 5 days before imaging observation. For lysosome live monitoring, SK-N-BE cells expressing LAMP1-eGFP protein are generated. Stable cell line is obtained upon plasmid transfection (epb-bsd-EIF1a-LAMP1eGFP) using Lipofec-tamine™ 2000 Transfection Reagent (ThermoFisher). The cells are then selected by Blasticidin (5 µg*/*mL) administration. For microtubule labelling, 1 µg*/*mL of Tubulin Tracker™ Deep Red (Invitrogen) is added to samples media and incubated for 30 min, followed by a washout before imaging and by changing the complemented media with RPMI 1640 Medium no phenol red (Gibco), to limit signal background during imaging observation. To perturbate lysosome motility, SK-N-BE cells expressing LAMP1-eGFP are treated with 10 µg*/*mL nocodazole (Merck) for 1 h, which destabilizes microtubules and, as described in [76], significantly increases the number of stationary lysosomes.

### Lysosome motion behavior segmentation

To differentiate between the “stop” and the “run” motion states of the lysosomes, we analyzed the residence time spent by the organelles in neighbouring positions during their movement. To estimate the residence time we divided the 3D space into a regular grid with a voxel size of 5 nm ×5 nm ×20 nm. Each voxel is then associated with a value which represents the number of trajectory points residing inside it. By multiplying with the trajectory sampling time we therefore obtain a measure of the residence time of the considered lysosome inside each voxel (Supplementary Fig. S8). We additionally smoothed the result with a 3D gaussian kernel to increase the contrast of spatial clusters of high residence time voxels and to reduce fluctuations.

Depending on the particular experiment, the optimal voxel residence time threshold may vary between 5 ms and 20 ms. All the trajectory points associated with a voxel with a value above the threshold are labelled as “stop”, while the others as “run”.

## Declarations

### Supplementary information

A supplementary file accompanies this article. The supplementary information contains theoretical details, additional methods, and additional figures.

## Acknowledgments

The authors thank all members of the Molecular Microscopy and Spectroscopy labs for the many helpful suggestions.

## Funding

This project has received funding from: the European Research Council, ”BrightEyes”, ERC-CoG No. 818699 (A.B., G.T., M.O.H., L.B. and G.V.); ”ASTRA”, ERC-SyG No. 855923 (I.B.); the European Union - Next Generation EU, PNRR MUR - M4C2 – Action 1.4 - Call “Potenziamento strutture di ricerca e creazione di “campioni nazionali di R&S” (CUP J33C22001130001), *National Center for Gene Therapy and Drugsbased on RNA Technolog* No. CN00000041 (I.B. and G.V.); by Associazione Italiana per la Ricerca Sul Cancro, ”Circular RNAs: novel players and biomarkers in tumorigenesis”, IG 2019 No. 23053 (I.B.); by Ministero dell’Istruzione, dell’Universitàe Della ricerca (MIUR), ”Non-coding RNAs, new players in gene expression regulation: studying their role in neuronal differentiation and in neurodegeneration”, PRIN2017 No. 2017P352Z4 (I.B.).

## Competing interests

G.V. has a personal financial interest (co-founders) in Genoa Instruments, Italy.

## Authors’ contributions

G.V. conceived the idea. G.V. designed the study. I.B., E.S., and G.V. supervised the project. A.B., G.T., and M.O.H. implemented the single-particle tracking architecture. A.B., G.T., M.O.H, and L.B., implemented the data acquisition and control system. A.B. and E.S. implemented the data analysis software. E.P., F.C., and I.B. designed the live-cell experiments. A.B., E.S., M.O.H. and G.V., analysed the data with the support of all other authors. A.B. and G.V. wrote the manuscript. All authors discussed the results and commented on the manuscript.

## Data availability

The experimental data generated and analysed in this study will be deposited in a publicly available Zenodo database.

**Code availability**

The Python data analysis source code and the Lab-view data-acquisition and control software used for the current study are available upon request.

## Notes

### Competing Interest Statement

Giuseppe Vicidomini has a personal financial interest (co-founders) in Genoa Instruments, Italy.

