## Supplementary Information for "4D Single-Particle Tracking with Asynchronous Read-Out SPAD-Array Detector"

### SUPPLEMENTARY INFORMATION NOTES

#### SI Note 1: Cramér-Rao bound derivation

In order to calculate the CRB, we adapt the mathematical framework described in [30] by introducing a structured detection. The first step requires us to model the measurement process. We assume that for each detector element $k$, both the signal and background countings follow a Poissonian distribution with mean $s_k$ and $b_k$ respectively. Furthermore, we assume the background to be independent of the emitter position $\mathbf{r}_\mathrm{e}$. Hence, the probability $p_k$ of detecting a photon in the k-th pixel can be written as

$$p_k(\mathbf{r}_\mathrm{e}) = \frac{s_k(\mathbf{r}_\mathrm{e}) + b_k}{\sum_{i=1}^{K} [s_i(\mathbf{r}_\mathrm{e}) + b_i]} \tag{1}$$

We define the SBR

$$SBR(\mathbf{r}_\mathrm{e}) = \frac{\sum_{k=1}^{K} s_k(\mathbf{r}_\mathrm{e})}{\sum_{k=1}^{K} b_k} \tag{2}$$

and insert it into Eq. 1 to obtain:

$$\begin{aligned} p_k(\mathbf{r}_\mathrm{e}) &= \frac{s_k(\mathbf{r}_\mathrm{e}) + b_k}{\sum_{i=1}^{K} s_i(\mathbf{r}_\mathrm{e}) + \sum_{i=1}^{K} s_i(\mathbf{r}_\mathrm{e})/SBR(\mathbf{r}_\mathrm{e})} = \\ &= \frac{s_k(\mathbf{r}_\mathrm{e})}{(1 + 1/SBR(\mathbf{r}_\mathrm{e})) \cdot \sum_{i=1}^{K} s_i(\mathbf{r}_\mathrm{e})} + \frac{b_k}{(1 + 1/SBR(\mathbf{r}_\mathrm{e})) \cdot \sum_{i=1}^{K} s_i(\mathbf{r}_\mathrm{e})} = \\ &= \frac{SBR(\mathbf{r}_\mathrm{e})}{SBR(\mathbf{r}_\mathrm{e}) + 1} \cdot \frac{s_k(\mathbf{r}_\mathrm{e})}{\sum_{i=1}^{K} s_i(\mathbf{r}_\mathrm{e})} + \frac{1}{SBR(\mathbf{r}_\mathrm{e}) + 1} \cdot \frac{b_k}{\sum_{i=1}^{K} b_i} \end{aligned} \tag{3}$$

If we assume the amount of signal photons to be proportional to the excitation intensity, we can substitute the ratios between photon countings in Eq. 3 with the ratios between the PSFs $h_k(\mathbf{r}_\mathrm{e})$ and obtain:

$$p_k(\mathbf{r}_\mathrm{e}) = \frac{SBR(\mathbf{r}_\mathrm{e})}{SBR(\mathbf{r}_\mathrm{e}) + 1} \cdot \frac{h_k(\mathbf{r}_\mathrm{e})}{\sum_{i=1}^{K} h_i(\mathbf{r}_\mathrm{e})} + \frac{1}{SBR(\mathbf{r}_\mathrm{e}) + 1} \cdot \frac{b_k}{\sum_{i=1}^{K} b_i} \tag{4}$$

The same operation applies also for Eq. 2:

$$\begin{aligned} SBR(\mathbf{r}_\mathrm{e}) &= SBR(\mathbf{0}) \cdot \frac{\sum_{k=1}^{K} h_k(\mathbf{r}_\mathrm{e})}{\sum_{k=1}^{K} h_k(\mathbf{0})} = \\ &= SBR_\mathrm{p} \cdot \frac{\sum_{k=1}^{K} h_k(\mathbf{r}_\mathrm{e})}{\sum_{k=1}^{K} h_k(\mathbf{0})} \end{aligned} \tag{5}$$

where $SBR_\mathrm{p} \equiv SBR(\mathbf{0})$ is the peak value of the SBR referred to a particle perfectly in focus.

With the model described by Eq. 4, we can compute the likelihood function $\mathcal{L}(\mathbf{r}_\mathrm{e}|\mathbf{n})$ considering a fluorophore in position $\mathbf{r}_\mathrm{e}$ providing $\mathbf{n} = (n_1, n_2, n_3, ..., n_K)$ photon counts for the $K = 25$ different PSFs as:

$$\mathcal{L}(\mathbf{r}_\mathrm{e}|\mathbf{n}) = \frac{N}{\prod_{k=1}^{K} n_k!} \cdot \prod_{k=1}^{K} p_k(\mathbf{r}_\mathrm{e})^{n_k} \tag{6}$$

where $N = \sum_{k=1}^{K} n_k$ is the total number of detected photons. We then use Eq. 6 to calculate the Fisher information matrix. For the 2D case $\mathbf{r}_\mathrm{e} = (x_\mathrm{e}, y_\mathrm{e})$ we obtain:

$$\begin{aligned} \mathbf{F}(\mathbf{r}_\mathrm{e}) &= -\mathbb{E}\left(\begin{bmatrix} \frac{\partial^2 \ln[\mathcal{L}(\mathbf{r}_\mathrm{e}|\mathbf{n})]}{\partial x^2} & \frac{\partial^2 \ln[\mathcal{L}(\mathbf{r}_\mathrm{e}|\mathbf{n})]}{\partial x \partial y} \\ \frac{\partial^2 \ln[\mathcal{L}(\mathbf{r}_\mathrm{e}|\mathbf{n})]}{\partial x \partial y} & \frac{\partial^2 \ln[\mathcal{L}(\mathbf{r}_\mathrm{e}|\mathbf{n})]}{\partial y^2} \end{bmatrix}\right) \\ &= N \cdot \sum_{k=1}^{K} \frac{1}{p_k} \cdot \begin{bmatrix} \left(\frac{\partial p_k(\mathbf{r}_\mathrm{e})}{\partial x}\right)^2 & \frac{\partial p_k(\mathbf{r}_\mathrm{e})}{\partial x}\frac{\partial p_k(\mathbf{r}_\mathrm{e})}{\partial y} \\ \frac{\partial p_k(\mathbf{r}_\mathrm{e})}{\partial y}\frac{\partial p_k(\mathbf{r}_\mathrm{e})}{\partial x} & \left(\frac{\partial p_k(\mathbf{r}_\mathrm{e})}{\partial y}\right)^2 \end{bmatrix} \end{aligned} \tag{7}$$

Finally, inverting the Fisher matrix we obtain the CRB as

$$\sigma_\mathrm{CRB}(\mathbf{r}_\mathrm{e}) = \sqrt{\frac{1}{2}\mathrm{Tr}(\mathbf{F}^{-1})} \tag{8}$$

To model a localization performed at a fixed acquisition time, we define the scalar parameter $N_\mathrm{p}$ as the peak number of detected photons for a particle perfectly in focus ($\mathbf{r}_\mathrm{e} = \mathbf{0}$). Thus, using Eq. 5, we can derive $N(\mathbf{r}_\mathrm{e})$ including the noise:

$$N(\mathbf{r}_\mathrm{e}) = N_\mathrm{p} \cdot \left(\frac{SBR_\mathrm{p}}{SBR_\mathrm{p}+1} \cdot \frac{\sum_{k=1}^{K} h_k(\mathbf{r}_\mathrm{e})}{\sum_{k=1}^{K} h_k(\mathbf{0})} + \frac{1}{SBR_\mathrm{p}+1}\right) \tag{9}$$

As pointed out in the main text, provided a fixed system overall PSF $h(\mathbf{r}, \mathbf{i})$, the parameters $SBR_\mathrm{p}$ and $N_\mathrm{p}$ contains all the information necessary to describe the localization experimental condition.

### SI Note 2: Localization estimators characterization

#### Maximum likelihood estimation

The MLE calculates the position of the fluorophore by maximizing the likelihood function already described in Eq. 6:

$$\hat{\mathbf{r}}_{\mathrm{e}} = \operatorname{argmax}\left(\mathcal{L}(\mathbf{r}_{\mathrm{e}}|\mathbf{n})\right) \tag{10}$$

Evidently, the estimation relies on the usage of a proper model of the PSF as well as the characterization of the noise represented by $SBR_{\mathrm{p}}$. The MLE experimentally provides an unbiased and linear localization in all directions, as shown in Fig. SN1.

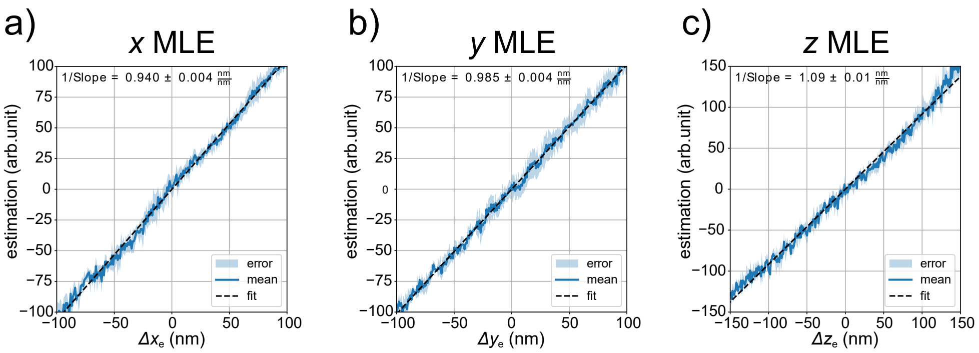


**SI Note Figure SN1 Experimental characterization of the maximum likelihood estimation in the three spatial dimensions. a-c)** Calibration curves obtained by moving with the piezoelectric stage a 40 nm fluorescent bead several times for each direction and localizing its position with the MLE ($SBR_{\mathrm{p}} = 50$).

#### Fast estimators

The high amount of calculations necessary to perform the estimation with the MLE makes its implementation in real-time unfeasible. We consequently define here a set of faster estimators, which allows us to gain computational speed at the expense of accuracy.

The information about any lateral displacements is contained in the position of the emission PSF in the microimage. Therefore, we can use the centroid estimator to obtain a fast lateral localization of the particle. Given a microimage $I(i, j) \equiv I(i, j \mid \mathbf{r}_{\mathrm{e}})$ and proper calibration factors $k_{\mathrm{x}}$ and $k_{\mathrm{y}}$ we define:

$$\hat{x}_\mathrm{e} = k_\mathrm{x} \cdot \frac{\sum\limits_{i=-2}^{2}\sum\limits_{j=-2}^{2} I(i,j)\cdot i}{\sum\limits_{i=-2}^{2}\sum\limits_{j=-2}^{2} I(i,j)}$$
$$\hat{y}_\mathrm{e} = k_\mathrm{y} \cdot \frac{\sum\limits_{j=-2}^{2}\sum\limits_{j=-2}^{2} I(i,j)\cdot j}{\sum\limits_{i=-2}^{2}\sum\limits_{j=-2}^{2} I(i,j)} \tag{11}$$

The centroid estimator correctly provides the position of the emission PSF as long as the intensity distribution in the microimage $I(i,j)$) is centrosymmetric with respect to the position to be estimated. This hypothesis holds in the absence of noise and when the emission is fully contained in the detector. When the position approaches the edge of the field-of-view (FoV) or noise is added, one or both of the requirements are not met and the performance is affected.

We use a simulated dataset to show the combination of the two parameters. As can be seen in Figs. SN2a,b, the first deviation from the ideal estimation happens because of cropping: the limited sFoV fails to image properly the emission PSF and the resulting microimage has a centroid shifted towards the center of the coordinate system. The effect of breaking the central symmetry is then worsened when flat noise is added. The recorded microimage can now be thought as the sum of the two independent patterns deriving from the signal and the background noise $I(i,j) = I_\mathrm{s}(i,j) + I_\mathrm{bkg}(i,j)$, each contributing to the centroid (Fig. SN2c). Mathematically:

$$\hat{x}_\mathrm{e} = k_\mathrm{x} \cdot \frac{\sum\limits_{i=-2}^{2}\sum\limits_{j=-2}^{2} [I_\mathrm{s}(i,j) + I_\mathrm{bkg}(i,j)]\cdot i}{\sum\limits_{i=-2}^{2}\sum\limits_{j=-2}^{2} [I_\mathrm{s}(i,j) + I_\mathrm{bkg}(i,j)]} =$$
$$= k_\mathrm{x} \cdot \frac{\sum\limits_{i=-2}^{2}\sum\limits_{j=-2}^{2} I_\mathrm{s}(i,j)\cdot i}{\sum\limits_{i=-2}^{2}\sum\limits_{j=-2}^{2} [I_\mathrm{s}(i,j) + I_\mathrm{bkg}(i,j)]} + k_\mathrm{x} \cdot \frac{\sum\limits_{i=-2}^{2}\sum\limits_{j=-2}^{2} I_\mathrm{bkg}(i,j)\cdot i}{\sum\limits_{i=-2}^{2}\sum\limits_{j=-2}^{2} [I_\mathrm{s}(i,j) + I_\mathrm{bkg}(i,j)]}$$

Defining $\widetilde{SBR} = \left(\sum_{i=-2}^{2}\sum_{j=-2}^{2} I_\mathrm{s}(i,j)\right) / \left(\sum_{i=-2}^{2}\sum_{j=-2}^{2} I_\mathrm{bkg}(i,j)\right)$ the SBR of the microimage and $\hat{x}_\mathrm{s}$, $\hat{x}_\mathrm{bkg}$ the centroid estimates of the signal and background patterns, we can finally rewrite:

$$\hat{x}_\mathrm{e} = \frac{\widetilde{\mathrm{SBR}}}{\widetilde{\mathrm{SBR}}+1}\cdot \hat{x}_\mathrm{s} + \frac{1}{\widetilde{\mathrm{SBR}}+1}\cdot \hat{x}_\mathrm{bkg} \tag{12}$$

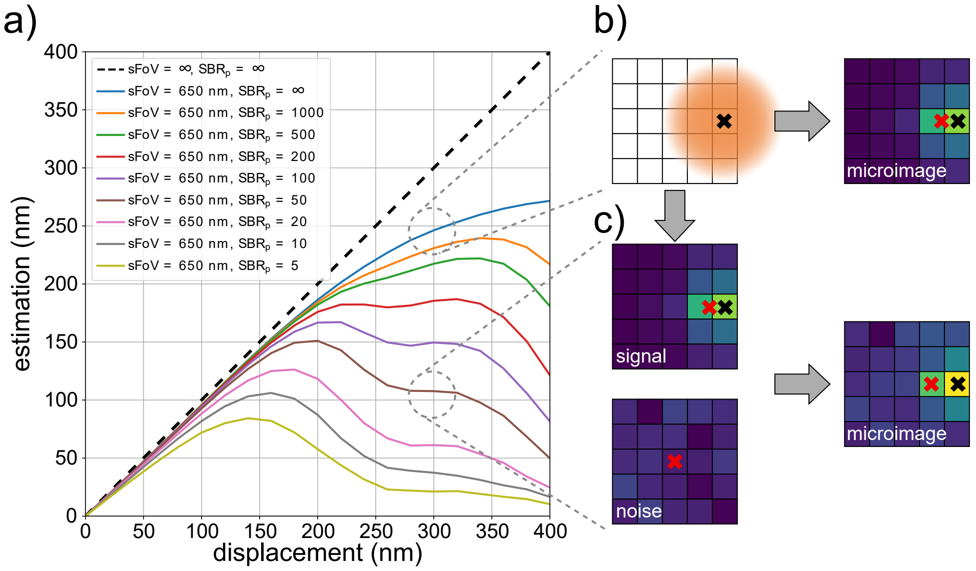


**SI Note Figure SN2 Simulated linearity of the centroid estimator as a function of the static field-of-view and the signal-to-noise ratio**. **a)** Simulated centroid estimation of a shift in the lateral direction. **b)** When the limited physical size of the detector is taken into account, the shifted emission PSF (orange spot) is cropped and the resulting microimage is centrosymmetric anymore. This leads to a difference between the estimated position $\hat{\mathbf{r}}_{\mathrm{e}}$ (red cross) and the actual one (black cross), with the former dragged towards the center of the sFoV. **c)** The presence of background noise ($SBR_{\mathrm{p}} = 50$) effectively creates another source of photons that adds up to the signal. The estimation of **b** is further modified by the centroid of this extra pattern and shows a bigger deviation from the ideal case.

A similar formula can be derived for $\hat{y}_{\mathrm{e}}$. Notably, the central symmetry of the emission PSF is preserved when the particle is shifted in the axial direction, making the lateral localization with the centroid independent from the axial position.

The latter information, on the contrary, is encoded in the shape the astigmatic emission PSF [82]. To leverage this characteristic, we define what we call the normalized difference estimator:

$$\begin{aligned}\hat{z}_{\mathrm{e}} = k_{\mathrm{z}} \cdot \frac{\sum_{i=-2}^{2} I(i,0) - \sum_{j=-2}^{2} I(0,j)}{\sum_{i=-2}^{2}\sum_{j=-2}^{2} I(i,j)} = \\ = k_{\mathrm{z}} \cdot \frac{s_{\mathrm{v}}(0) - s_{\mathrm{h}}(0)}{\sum_{i=-2}^{2}\sum_{j=-2}^{2} I(i,j)}\end{aligned} \tag{13}$$

where $k_z$ is a calibration factor and $s_v(j)$ and $s_h(i)$ are the sums performed along the j-th column and i-th row.

Intuitively, the direction along which the PSF is more elongated is the one containing the biggest fraction of the intensity. Assuming the PSF is placed at the coordinates $(i_p, j_p)$ and is horizontally stretched, then the sum of the intensity values along its main horizontal axis $s_h(i_p)$ is bigger than the sum along the main vertical axis $s_v(j_p)$. Since this is a pure geometrical effect, the division by the overall intensity is necessary to normalize the difference. To gain computational speed, the position of the main axes of the PSF is not estimated and the sum is arbitrarily performed over the central row and central column of the detector ($s_h(0)$ and $s_v(0)$). As a consequence, the estimator is linear and unbiased only when the emission is centered in the microimage and bias arises with increasing lateral displacements. As can be seen in Fig. SN3, a shift in the $y_e$ direction leads to a positive bias while a shift in the $x_e$ direction leads to a negative bias.

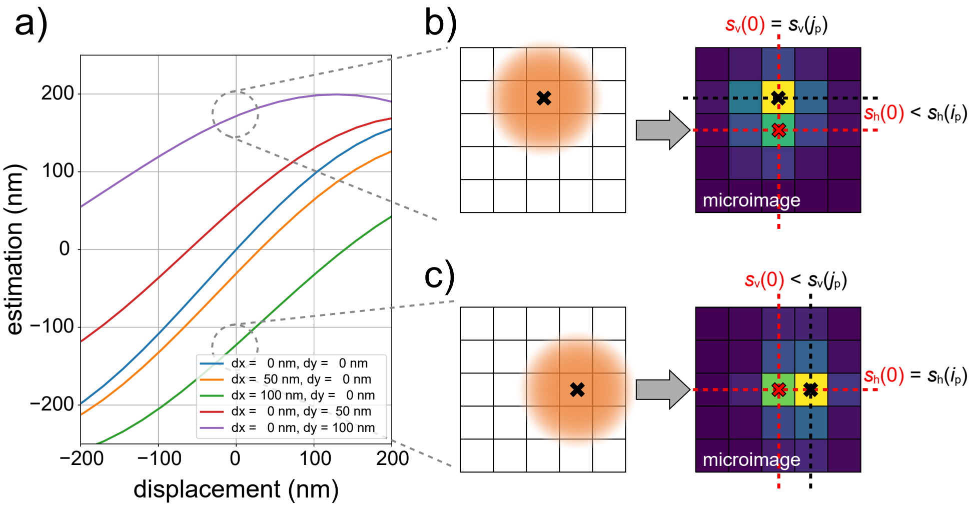


**SI Note Figure SN3 Simulated bias and linearity of the normalized difference estimator as a function of the lateral displacement. a)** Simulated axial estimation curves of the normalized difference estimator with increasing shifts in both the lateral directions ($dx$ and $dy$). **b)** When the emission is shifted in the vertical lateral direction (in this case $dy = 100\,\text{nm}$), the sum of the intensity in the central row $s_h(0)$ (red) is smaller than the sum of the intensity along the main horizontal axis of the emission PSF $s_h(i_p)$ (black). The estimation $\hat{z}_e \propto s_v(0) - s_h(0)$ is therefore biased towards the positive axial direction. **c)** A shift in the lateral horizontal direction ($dx = 100\,\text{nm}$) causes an analog situation as depicted in **b**. The sum along the central column $s_v(0)$ is smaller than the one of the main vertical axis $s_v(j_p)$ and the estimation $\hat{z}_e$ is consequently biased towards the negative axial direction.

In theory, the convergence of the feedback loop is threatened by a biased axial estimation, but this negative outcome is, in practice, counterbalanced by two main effects. First of all, the tracking procedure in the lateral direction keeps the sFoV locked on top of the particle and compensates for any lateral shift. Secondly, assuming the particle is in focus at the beginning of each estimation window, its intensity decreases while drifting laterally, according to the excitation PSF. Therefore the emission from peripheral positions contributes less to the microimage and the apparent position of the integrated intensity distribution is dragged towards the center of the detector.

In case the experiment requires it, it is nevertheless a good idea to make the axial estimation slower than the lateral one, improving axial accuracy at the cost of time resolution.

Despite all this possible sources of non-idealities, the centroid and normalized difference experimentally performs linearly in a region of hundreds of nanometers as shown in Fig. SN4.

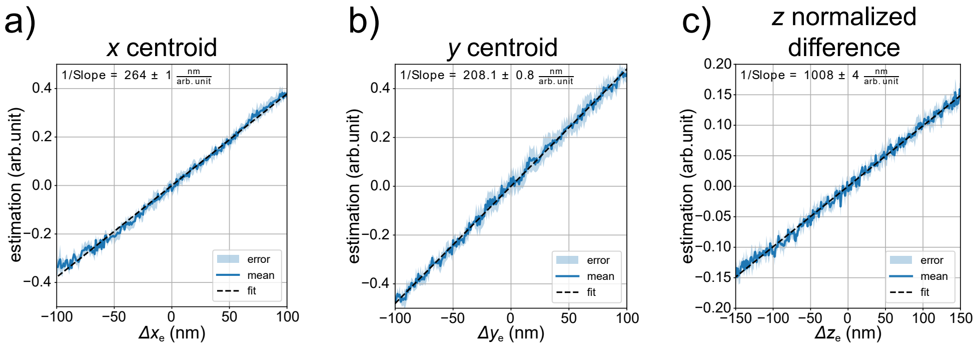


**SI Note Figure SN4 Experimental characterization of the centroid and normalized differential estimators. a-c)** The localization is performed laterally with the centroid and axially with the normalized difference estimators on the same dataset of Fig. SN1.

### SI Note 3: Statistical lifetime-based segmentation of the lysosome motion pattern

The correlation between the fluorescence lifetime and the motion state of lysosomes has potential implications for studying their diffusion behaviors. In fact, by analyzing the value of fluorescence lifetime, it may be possible to segment and study the motion of lysosomes in a simple and effective way. To test this hypothesis, we collect a set of 15 independent 4D tracking experiments and analyze them together.

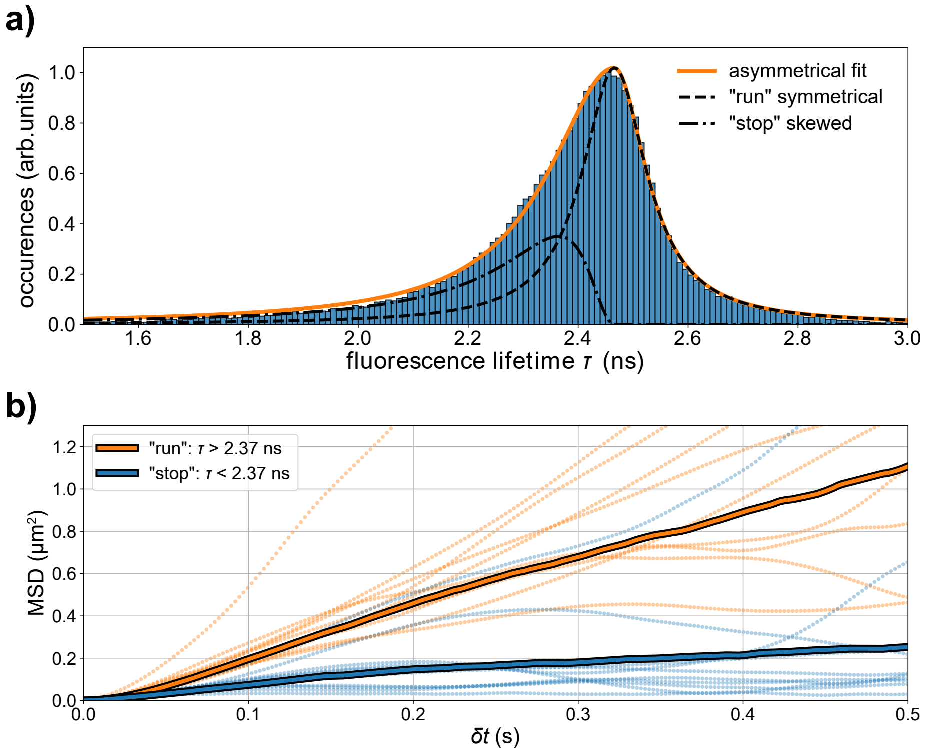


**SI Note Figure SN5 Segmentation of the lysosome state by the fluorescence lifetime. a)** Histogram of the fluorescence lifetime values measured in 15 independent RT-4D-SPT experiments on lysosomes. The shape is fitted with a custom asymmetrical Cauchy-Lorentz distribution for separating the contributions of the two motion states. The optimal segmentation threshold $\tau_{\mathrm{th}} \approx 2.37\,\mathrm{ns}$ is marked by the red line. **b)** mean squared displacement plot of the lysosome trajectories segmented with the fluorescence lifetime. The solid line curve is obtained by averaging over all the segments of all the 15 experiments (206 segments for the “run” state and 111 segments for the “stop” state). The small dots shows the MSD curves of 10 randomly selected trajectory segments for each motion state. The re-centering is performed at a fixed dwell time $\Delta t_{\mathrm{rc}}^{\mathrm{all}} = 2.5\,\mathrm{ms}$ and the lifetime is measured every $\Delta t_{\tau} = 10\,\mathrm{ms}$.

As depicted in Fig. SN5a, the histogram of all the measurements reveals a negatively skewed distribution of the fluorescence lifetime, which is consistent with the decrease in lifetime during the "stop" state previously reported in the main text. To determine an optimal threshold value for the segmentation of the trajectories, it is necessary to separate the two contributions. In the absence of a physical model that would allow us to derive a theoretical fitting function, we define a custom skewed curve based on the Cauchy-Lorentz distribution:

$$f(\tau; k, \mu, \gamma_r, \gamma_l) = \begin{cases} k \cdot \frac{\gamma_r}{\gamma_r^2 + (\tau-\mu)^2} & \text{if } \tau > \mu \\ \frac{k}{\gamma_r} \cdot \frac{\gamma_l^2}{\gamma_l^2 + (\tau-\mu)^2} & \text{if } \tau < \mu \end{cases} \tag{14}$$

where $\mu$ is the mode of the distribution, $\gamma_r$ and $\gamma_l$ the right and left half width at half maximum and $k$ is a scaling factor. The fitted curve agrees well with the experimental data, yielding the parameters $\mu = 2.466 \pm 0.001$ ns, $\gamma_r = 0.071 \pm 0.001$ ns, and $\gamma_l = 0.146 \pm 0.001$ ns.

Notably, the peak of $f$ is in agreement with the expected fluorescence lifetime for the GFP. If we assume that the "run" state generates a symmetrical Lorentzian distribution centered at $\mu$, we can divide the custom function into two distinct contributions $f = f_{run} + f_{stop}$:

$$f_{run}(\tau; k, \mu, \gamma_r) = k \cdot \frac{\gamma_r}{\gamma_r^2 + (\tau - \mu)^2} \tag{15}$$

$$f_{stop}(\tau; k, \mu, \gamma_r, \gamma_l) = \begin{cases} 0 & \text{if } \tau > \mu \\ \frac{k}{\gamma_r} \cdot \frac{(\tau-\mu)^2 \cdot (\gamma_l^2 - \gamma_r^2)}{(\gamma_l^2 + (\tau-\mu)^2) \cdot (\gamma_r^2 + (\tau-\mu)^2)} & \text{if } \tau < \mu \end{cases} \tag{16}$$

Assuming $f_{run}$ and $f_{stop}$ are the probability distributions of measuring a certain $\tau$ in a given motion state, we determine the optimal threshold as the point in which both processes are equally likely $\tau_{th} \approx 2.37$ ns. Trajectory points with a lifetime below $\tau_{th}$ are assigned the "stop" state and the others the "run" state.

We expect here the segments labeled as "run" state to display a faster movement compared to those labeled as "stop" state. To confirm this expectation, we compute the MSD of the two populations (Fig. SN5b). The analysis reveals that the "run" state population indeed exhibits faster diffusion. Contrary to the expectation, the linearity of the curve suggests free diffusion. Nevertheless, we believe that the free diffusion motion is just an artifact caused by averaging all the "run" states of all the vesicles together. In fact, each MSD curve of single "run" state segments, exhibit a different non-brownian diffusion, thus reflecting the variability of tasks performed by the lysosomes. On the other hand, the "stop" state curve shows a strong sub-diffusion behavior, which is an indication of confinement of the lysosomes.

### SUPPLEMENTARY INFORMATION FIGURES

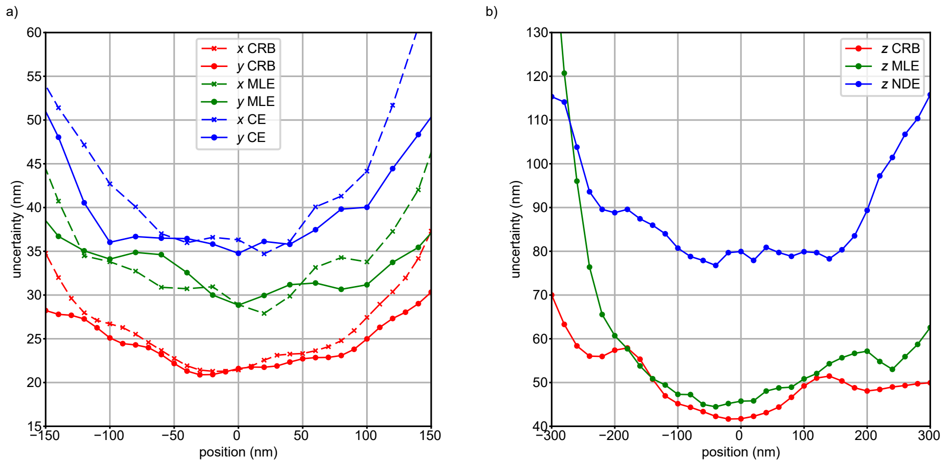


**SI Figure S1 Line profiles of the planar localization uncertainty maps**. **a)** Line profiles of Fig. 2a-c along the $x$ and $y$ directions. **b)** Line profiles of Fig. 2d-f along the $z$ direction.

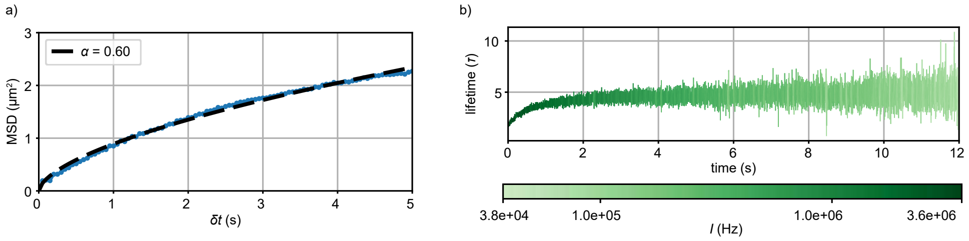


**SI Figure S2 Time evolution of the trajectory of Fig. 3b-d. a)** MSD of the trajectory with fitting showing anomalous subdiffusion ($\alpha < 1$). **b)** 2D plot of the time evolution of the fluorescence lifetime $\tau$ with color used to visualize the photon flux.

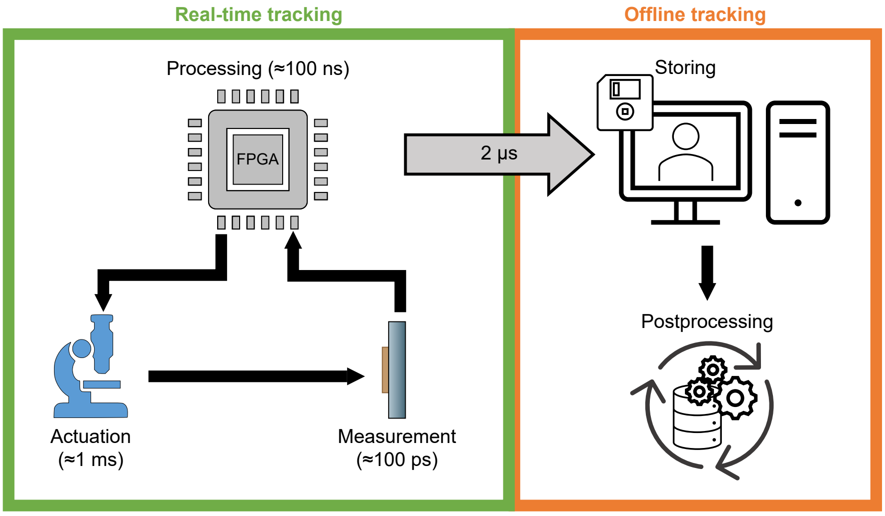


**SI Figure S3 Overview of the main stages of the tracking procedure**. Our tracking approach features both a real-time feedback loop and an offline part. The feedback loop is composed of three cyclic steps: measurement, processing and actuation, with the latter currently representing the real-time speed bottleneck. The offline part relies on saved data and can be used to postprocess the trajectories to achieve higher temporal resolution ("rebinning") and lower spatial uncertainty ("refining").

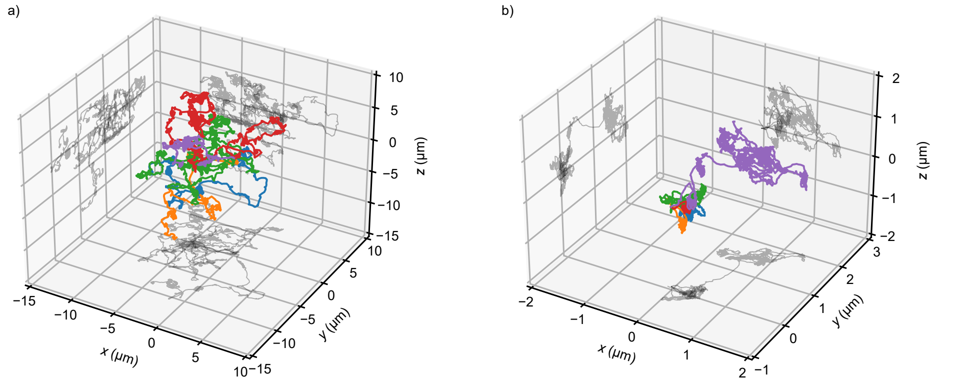


**SI Figure S4  3D trajectories of the lysosomes before and after the addition of nocodazole**. **a)** 3D plot of 5 example trajectories of lysosomes moving inside living human SK-N-BE cells. **b)** 3D plot of 5 example trajectories of lysosomes moving in the same sample of **a** after the addition of nocodazole. All the measurements are performed with a fixed re-centering time $\Delta t_{\mathrm{rc}}^{\mathrm{all}} = 2.5\,\mathrm{ms}$.

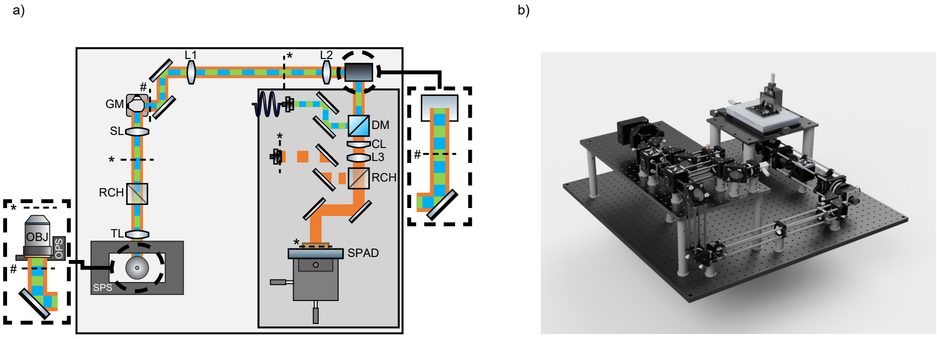


**SI Figure S5 Microscope setup**. **a)** 2D sketch of the optical beam path. OBJ = objective, SPS = sample piezoelectric stage, OPS = objective piezoelectric stage, TL = tube lens, SL = scan lens, GM = galvanometric mirrors, DM = dichroic mirror, L1 = 200 mm, L2 = 150 mm, L3 = 300 mm, CL = cylindrical lens (f = 1000 mm), RCH = removable cube holder, * = imaging plane, # = phase plane. **b)** 3D render of the microscope.

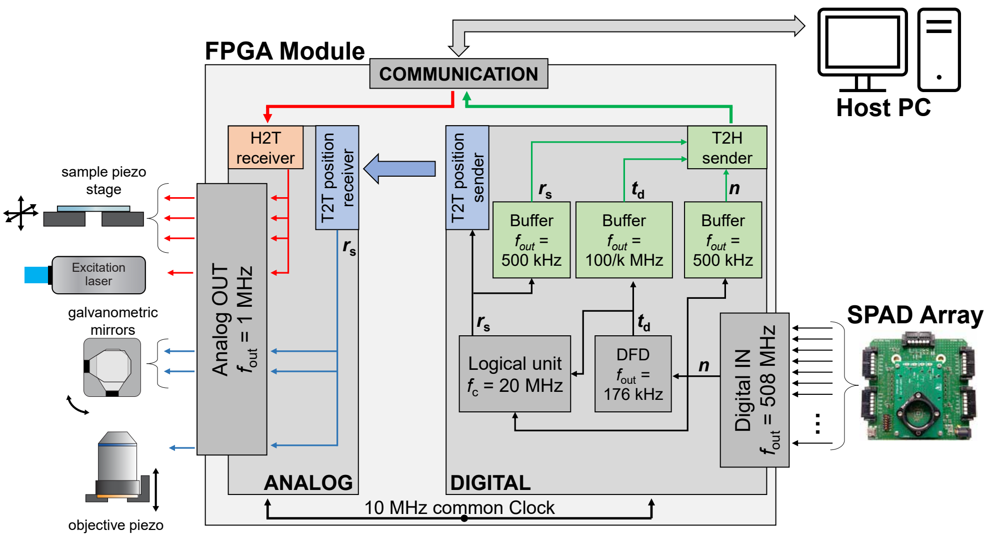


**SI Figure S6 Control firmware for tracking**. The two FPGA cards are fed a common clock at 10 MHz from which all the other clocks are derived. Each functional block within the system operates at a specific output frequency $f_{\text{out}}$, indicating the rate at which its output becomes available. The logical unit, exceptionally, is characterized by its internal cycle frequency $f_c$, which denotes the fixed clock at which the logical steps are executed. In the schematic representation, the color green is used to highlight the lines and blocks involved in transferring the experimental data from the digital target to the host PC (T2H). Conversely, the color red is used to indicate control commands sent from the host PC to the analog target (H2T) relating to the piezoelectric stage positioning and laser power. The blue pathway represents the internal transfer of the scanning position from the digital card to the analog card (T2T). This communication utilizes the eight trigger lines and employs a custom-written protocol to ensure an instruction communication rate of at least $\approx$5 MHz. $\mathbf{n}$ = photon counting registers, $\mathbf{t}_{\text{d}}$ = time of detection of the photons and the pulsed excitation, $\mathbf{r}_{\text{s}}$ = scanning position.

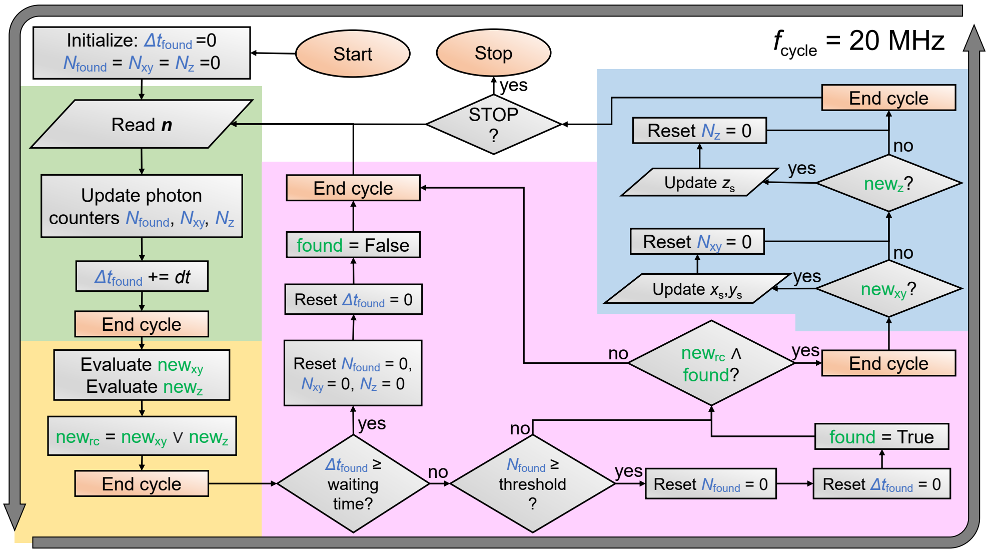


**SI Figure S7 Flowchart diagram of the tracking logical unit**. The decision-making algorithm consists of four main steps, each represented by a different background color: green for "collection," yellow for "evaluation," pink for "decision," and blue for "re-centering." These steps are executed at a clock frequency of 20 MHz and are visually ended by an orange cycle breaking block. The "decision" step serves as a crucial junction in the algorithm, incorporating the necessary logical decisions to manage the real-time response of the system. It can lead to the subsequent "re-centering" step or loop back to the "collection" step. The algorithm utilizes both boolean variables (highlighted in green) and numeric variables (highlighted in blue). In the current implementation, it only takes the photon counting registers **n** as input. By following this algorithmic flow, the system can effectively collect and evaluate data, make informed decisions, and perform re-centering actions as needed.

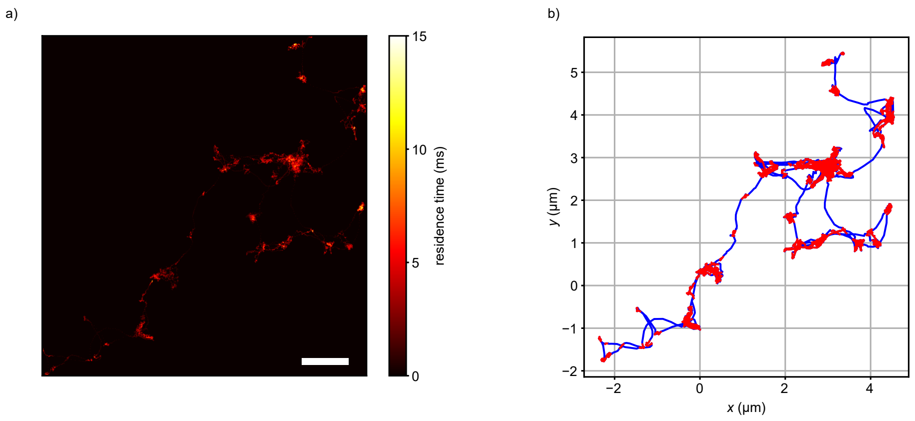


**SI Figure S8 Lysosome residence time and motion state segmentation**. **a)** 2D maximum projection of the voxel residence time for the lysosome trajectory of Fig. 4. Voxel size is 5 nm ×5 nm ×20 nm. Scalebar = 1 µm. **b)** 2D result of the segmentation of **a** with a threshold of 5 ms.

### SUPPLEMENTARY INFORMATION TABLES

| Diameter | $\lambda_{exc}$ | Part number | Figure reference |
|---|---|---|---|
| 20 nm | 561 nm | Invitrogen™ FluoSpheres™ F8786 | 2, S1 |
| 40 nm | 561 nm | Invitrogen™ FluoSpheres™ F8794 | 3 |
| 100 nm | 488 nm | Invitrogen™ FluoSpheres™ F8803 | 3, S2 |

**SI Table ST1 List of all the fluorescent beads utilized in the various experiments.**
